# Fine-scale spatial genetic structure in a locally abundant native bunchgrass (*Achnatherum thurberianum*) including distinct lineages revealed within seed transfer zones

**DOI:** 10.1101/2022.06.22.497217

**Authors:** Carolina Osuna-Mascaró, Alison C. Agneray, Lanie M. Galland, Elizabeth A. Leger, Thomas L. Parchman

## Abstract

Analyses of the factors shaping spatial genetic structure in widespread plant species are important for understanding evolutionary history and local adaptation and have applied significance for guiding conservation and restoration decisions. Thurber’s needlegrass (*Achnatherum thurberianum*) is a widespread, locally abundant grass that inhabits heterogeneous arid environments of western North America and is of restoration significance. It is a common component of shrubland steppe communities in the Great Basin Desert, where drought, fire, and invasive grasses have degraded natural communities. Using a reduced representation sequencing approach, we generated SNP data at 5,677 loci across 246 individuals from 17 *A. thurberianum* populations spanning five previously delineated seed zones from the western Great Basin. Analyses revealed pronounced population genetic structure, with individuals forming consistent geographical clusters across a variety of population genetic analyses and spatial scales. Low levels of genetic diversity within populations, as well as high population estimates of linkage disequilibrium and inbreeding, were consistent with self-fertilization as a contributor to population differentiation. Moreover, variance partitioning and partial RDA indicated local adaptation to the environment as an additional factor influencing the spatial distribution of genetic variation. The environmental variables driving these results were similar to those implicated in recent genecological work which inferred local adaptation in order to delineate seed zones. However, our analyses also reveal a complex evolutionary history of *A. thurberanium* in the Great Basin, where previously delineated seed zones contain distantly related populations. Overall, our results indicate that numerous factors shape genetic variation in *A. thurberianum* and that evolutionary history, along with differentiation across distinct geographic and environmental scales, should be considered for conservation and restoration plans.

## Introduction

Identifying the factors that drive patterns of genetic variation among plant populations is important for understanding ecological and evolutionary processes and has applied significance for conservation and ecological restoration (Sork et al., 1999; Hedrick, 2005; Holderegger and Wagner 2008; Sommer et al., 2013). The spatial distribution of genetic variation reflects evolutionary processes, including drift, migration, and selection, which shape the standing variation and the evolutionary potential of populations. Therefore, quantifying spatial genetic structure and the factors shaping it can help assess the degree of population connectivity, the scale of and potential for local adaptation to environmental variation, and, consequently, the persistence of plant populations faced with environmental change (Bauert et al., 1998; Booy et al., 2000; Manel et al., 2003). Such analyses can also be used to guide conservation and restoration decisions using biologically meaningful information (Ottewell et al., 2016; Carvalho et al., 2021). During the last decade, high throughput sequencing approaches have substantially improved our ability to quantify spatial genetic structure and infer its causes across populations of ecologically significant non-model organisms (Andrews et al., 2016; Breed et al., 2019; Hohenlohe et al., 2021).

For plant species with large distributions spanning heterogeneous environments, spatial genetic structure can be shaped by numerous factors, including geological, historical, and environmental factors, as well as life-history variation (Holderegger et al., 2010). Across large geographic scales, genetic differentiation among populations can be expected as gene flow decays with increasing geographic distance and across geological barriers, commonly resulting in isolation by distance (Wright, 1943; Gavrilets et al., 2000; Hoskin et al., 2005). However, environmental and ecological factors may also play a role in shaping spatial genetic structure (Alvarez et al., 2009; Storfer et al., 2010; Paz et al., 2015; Mosca et al., 2018). Environmental variation can directly influence genetic differentiation by causing local adaptation and indirectly by generating Isolation by Environment (IBE; Shafer and Wolf, 2013; Wang and Bradburd, 2014), where gene flow is reduced across environmental gradients or selection against migrants occurs (Kawecki and Ebert, 2004). Thus, strong population genetic differentiation can occur across regions experiencing different ecological and environmental conditions (Ortego et al., 2012; Orsini et al., 2013; Wang et al., 2013; Wang and Bradburd, 2014).

Mating system also influences patterns of population genetic structure in plants (Williams et al., 2001; Duminil et al., 2007; Gamba and Muchhala, 2020), due to variation in the frequency with which offspring are produced asexually, through self-fertilization, or via sexual outcrossing (Holsinger, 2000). Compared to outcrossers, asexual and selfing plants often have reduced levels of within-population genetic diversity. In particular, selfing plants often exhibit low population genetic variation and high inbreeding which can lead to relatively pronounced patterns of population differentiation (Hamrick and Godt, 1996; Volis et al., 2010; Wagner et al., 2012; Huang et al., 2021). Thus, selfing plants may show stronger patterns of population structure at a local scale and lower genetic diversity than outcrossing plants with higher genetic diversity and non-structured populations. Mating system could thus impact another decision increasingly common in restoration, which is whether to combine collections from multiple populations, either to deliberately increase diversity or as a practical decision when there are simply not enough seeds available from one population (St. Clair et al., 2020). While there is concern about the potential for outbreeding depression when combining populations of outbreeding species (Templeton, 1986; Hufford et al., 2012), these concerns could be reduced for highly inbreeding species.

In complex environments, isolation and convergent evolution can result in similar but independently derived phenotypes in populations with very different evolutionary histories (St. Clair et al., 2013; Massatti et al., 2018). While common in natural systems, these patterns may confound restoration efforts, leading to situations where practitioners may be choosing between seed sources that either possess phenotypes that are likely to be adaptive in a given site but are distantly related to plants that used to occur there, or are more closely related to former inhabitants, maintaining historical patterns of gene flow, but with sub-optimal phenotypes that may not survive as well in restoration sites (Leger et al., 2021). Historically, evolutionary history has been commonly considered in conservation planning for rare species (Verdú et al., 2012), while an emphasis on adaptive phenotypes has been the focus for delineating seed transfer zones for restoration of widely-distributed species (Baughman et al., 2019; Pedlar et al., 2021). However, due to advances in DNA sequencing technology, it has recently become possible to consider evolutionary history as well as both genomic and phenotypic evidence for local adaptation for widely distributed species (Massatti et al., 2018; Breed et al., 2019). Here we consider how evolutionary history and environmental variation shape landscape genetic structure in a widespread grass species of restoration significance in the Great Basin Desert, for which seed transfer zones have been previously inferred based on phenotypic evidence for local adaptation (Johnson et al. 2017).

The Great Basin is the most extensive cold desert in North America, with an area of about 540,000 km^2^, that harbors significant environmental heterogeneity and biological diversity, including high levels of genetic diversity and local adaptation across extensive environmental gradients created by the complex, repeated basin and range topography (West, 1983; Pilliod et al., 2017; Baughman et al., 2019; Faske et al., 2021). However, in recent decades, the combined effects of altered fire regimes, invasive annual grasses, and human land use have led to widespread degradation and fragmentation of habitats in the Great Basin (Balch et al., 2013; Pilliod et al., 2013). In particular, cheatgrass (*Bromus tectorum*) and other non-native annual species have transformed native shrublands into invasive-dominated grasslands (Knapp, 1996; Parkinson et al., 2013; Nagy et al., 2021). These factors, along with climate change, are contributing to native plant species declines, especially native grasses, many of which are decreasing in abundance (Chambers and Wisdom, 2009; Svejcar et al., 2017). Overall, restoration efforts are increasing in response to global initiatives (Dudley et al., 2020; Stange et al., 2021), especially in drylands such as the Great Basin (Pilliod et al., 2017; Shackelford et al., 2021). Despite the widely recognized importance of considering spatial genetic structure and local adaptation for restoration planning (Knapp and Rice, 1994; Hufford and Mazer, 2003; McKay et al., 2005; Breed et al., 2019), and the growing number of plants in this region with phenotype-based seed transfer zones (TRM Seed Zone Applications: https://www.fs.fed.us/wwetac/threat-map/TRMSeedZoneMapper.php), we lack population genomic perspectives for most Great Basin native plants of restoration significance (but see Massatti et al., 2018).

Among the Great Basin grass species of restoration interest is Thurber’s needlegrass, *Achnatherum thurberianum* (Piper) Barkworth, a widespread perennial bunchgrass species and an essential component of many sagebrush communities (Johnson et al., 2017). *Achnatherum* (Poaceae, subfamily Pooideae, tribe Stipeae) consists of large perennial grasses that grow in temperate grassland and savannah habitats (Soreng et al., 2015; Soreng et al., 2017), many of which are thought to self-fertilize (Jones and Nielson, 1989; Durka et al., 2013; Kraehmer, 2019). Genetic boundaries of *Achnatherum* are controversial (Jacobs et al., 2007; Cialdella et al., 2010; Romaschenko et al., 2010, 2012; Peterson and Romaschenko, 2019), and accordingly, the taxonomy of the genus has been reviewed several times (Hamasha et al., 2012). In particular, *A. thurberianum* has been previously classified as *Stipa thurberiana* (Piper), (Circ. Div. Agrostol. U.S.D.A., 1900), *A. thurberianum* (Piper), (Barkworth, 1993), and the more recent although not yet widely in use *Eriocoma thurberiana* (Piper) (Peterson and Romaschenko, 2019; see the Missouri Botanical Garden’s taxonomic database; http://www.tropicos.org). Studies on related species in Eurasia (from the tribe Stipeae) based on traditional molecular markers indicated pronounced population differentiation and low diversity consistent with a selfing mating system and demographic processes shaping population differentiation at small spatial scales (Wagner et al., 2012; Durka et al., 2013). Other studies on Eurasian species have indicated the potential for climate to shape local adaptation and population genetic structure (Hamasha et al., 2013; Gao et al., 2018; Schubert et al., 2019). Recent genecological work on *A. thurberianum* phenotypes across the Great Basin illustrated local adaptation in response to temperature and precipitation variation, which led to the formation of seed transfer zones for this species (Johnson et al., 2017). Specifically, in that study, populations from warmer and drier regions generally exhibited earlier flowering time and narrower leaves than those from cooler wetter regions (Johnson et al., 2017). However, we currently lack a baseline perspective on the spatial distribution of genetic variation for *A. thurberanum* in the Great Basin region, which is important because a complex evolutionary history could have unintended consequences for seed sourcing if transfer zones are delineated based on only climate, proximity, or phenotypic evidence for local adaptation (Massatti et al., 2018). Analyses of evolutionary history and how landscape genetic variation is shaped by environmental, geographic, and life history variation stand to improve understanding of the biology of this widespread species and provide a critical context for understanding its evolution and restoration potential, including the application of phenotype-based seed transfer zones (Johnson et al., 2017).

Here, we used reduced representation sequencing (ddRADseq) to characterize spatial genetic variation across 17 *A. thurberianum* populations distributed across the southwestern Great Basin, in an area representing five of the twelve seed zones proposed by Johnson et al. (2017). We examined phylogeographic patterns and spatial genetic structure across populations and considered how geographical and environmental factors influenced them. We also quantified levels of genetic diversity, linkage disequilibrium, and inbreeding coefficients within populations to evaluate the extent to which mating system might influence standing variation and fine-scale differentiation across the sampled populations and asked to what degree existing seed zones reflected evolutionary history. We expected to see pronounced regional and fine-scale spatial genetic structure, influenced by both mating system and local adaptation to environmental variation in this widespread grass. Further, we expected that we might find evidence of multiple lineages represented within seed zones, given the complexity of the basin and range topography and the potential for convergent evolution during the process of local adaptation.

## Material and methods

### Plant material

We collected leaf material from 17 localities in the western Great Basin during the Fall of 2017 from a range of 20 to 39 plants per location (Figures 1a, 2a; Table 1). We sampled these locations because they hosted multiple native species that could potentially serve as restoration seed sources for this region; collections of other species from these locations are being used for additional restoration genetic studies (e.g., Faske et al., 2021; Agneray et al., 2022). In addition, these localities spanned five seed zones proposed by Johnson et al., (2017). Means for representative environmental variables for each population were obtained as described below and can be found in Suppl. Table 1. Because of the complex topography in this region, note that proximate populations are not necessarily within the same seed zone (Figure 1a). A total of 246 individuals, from a range of 12 to 18 plants per location, were included in analyses after DNA extraction and quality screening (see below).

**Figure 1.**
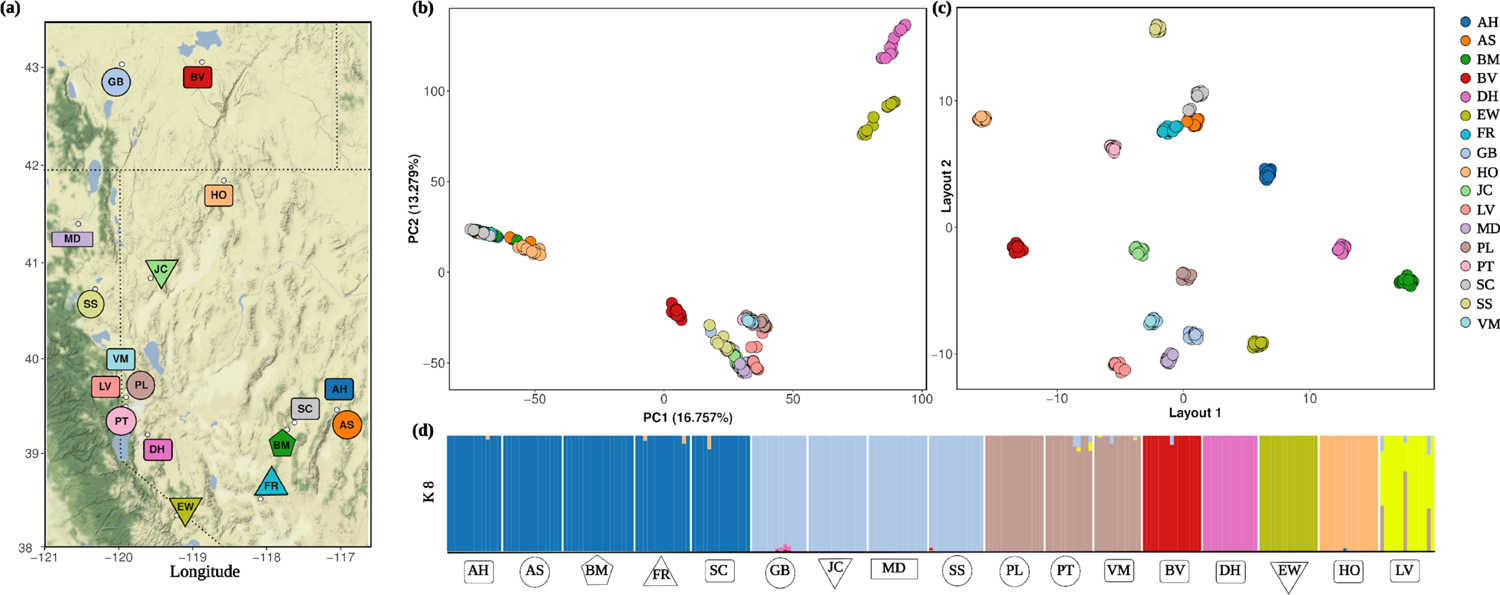
Population genetic structure of 17 populations of *Achnatherum thurberianum* based on 5,677 SNPs. a) Map of the sampled locations with each population code. Each population is colored consistently in panels a, b, and c, and represented by one of five shapes corresponding to the seed zone of Johnson et al. (2017) containing each population (Table 1). b) The first and second principal components (PCs) resulting from PCA on the genotypic data plotted for each individual. c) Each individual plotted for the first two axes from a Uniform Manifold Approximation and Projection (UMAP) clustering analysis based on the genotypic data. d) ADMIXTURE plot representing estimated ancestry coefficients for each individual, following results of an analysis with K set to eight ancestral populations.

**Figure 2.**
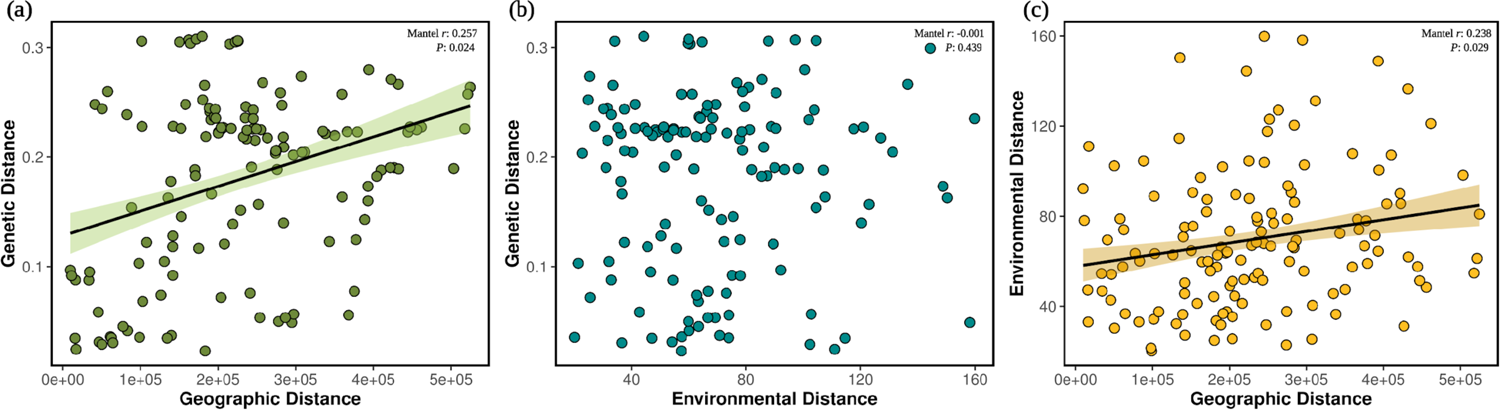
Mantel test results and relationships between genetic (Nei’s D) and geographic distances (km) (a), genetic (Nei’s D) and environmental distances (b), and geographic (km) and environmental distances (c).

**Table 1.** Population genetic summary statistics for each sampled *A. thurberianum* population. For each sampling location (population), the table shows the number of individuals (N), expected (H_e_) and observed heterozygosity (H_o_), inbreeding coefficient (F_IS_), nucleotide diversity (π), Tajima’s D (D), standardized index of association (r^—^_d_), and index of association (I_A_). Significant F_IS_ results based on bootstrap coefficient intervals are depicted with an asterisk. The seed zone for each population corresponds to those proposed by Johnson et al., (2017). The MD population has no seed zone information as it was located outside of the Johnson et al., (2017) projected area.

| Population name,<br>State | Population<br>abbr. | N | $H_e$ | $H_o$ | $F_{IS}$ | $\pi$ | $D$ | $r_d^-$ | $I_A$ | Seed<br>zone |
| --- | --- | --- | --- | --- | --- | --- | --- | --- | --- | --- |
| Austin Hwy, NV | AH | 14 | 0.029 | 0.007 | 0.719* | 0.039 | 0.487 | 0.283 | 122.350 | 8 |
| Austin Summit, NV | AS | 15 | 0.028 | 0.010 | 0.649* | 0.027 | 0.519 | 0.226 | 93.120 | 5 |
| Bald Mountain, NV | BM | 18 | 0.025 | 0.008 | 0.671* | 0.025 | 1.030 | 0.186 | 58.549 | 4 |
| Buena Vista, OR | BV | 15 | 0.066 | 0.029 | 0.509* | 0.067 | 0.339 | 0.019 | 21.443 | 8 |
| Dayton Hill, NV | DH | 14 | 0.044 | 0.011 | 0.728 | 0.045 | 0.577 | 0.137 | 87.699 | 8 |
| East Walker, CA | EW | 15 | 0.028 | 0.011 | 0.704 | 0.028 | 0.307 | 0.314 | 148.414 | 11 |
| Finger Rock, NV | FR | 14 | 0.026 | 0.010 | 0.675* | 0.026 | 0.487 | 0.271 | 106.050 | 7 |
| Grey's Butte, NV | GB | 14 | 0.060 | 0.024 | 0.522* | 0.061 | 0.311 | 0.090 | 89.900 | 5 |
| Hwy 140, NV | HO | 15 | 0.052 | 0.024 | 0.484* | 0.052 | 0.252 | 0.025 | 22.540 | 8 |
| Jones Canyon, NV | JC | 15 | 0.059 | 0.014 | 0.740* | 0.058 | 0.752 | 0.102 | 86.020 | 11 |
| Long Valley, CA | LV | 14 | 0.075 | 0.012 | 0.816* | 0.074 | 0.512 | 0.236 | 270.389 | 8 |
| Modoc, CA | MD | 15 | 0.058 | 0.016 | 0.715* | 0.057 | 0.483 | 0.047 | 43.020 | NA |
| Peavine Low, NV | PL | 15 | 0.049 | 0.032 | 0.368* | 0.054 | 0.787 | 0.180 | 121.440 | 5 |
| Patagonia, NV | PT | 12 | 0.035 | 0.024 | 0.344* | 0.041 | 0.183 | 0.133 | 81.633 | 5 |
| Smith Creek, NV | SC | 15 | 0.030 | 0.010 | 0.611 | 0.029 | 0.536 | 0.329 | 143.590 | 8 |
| Spanish Springs, CA | SS | 14 | 0.060 | 0.024 | 0.551 | 0.067 | 0.273 | 0.029 | 29.786 | 5 |
| Virginia Mountains, NV | VM | 12 | 0.049 | 0.017 | 0.603 | 0.054 | 0.401 | 0.084 | 64.681 | 8 |

### Library preparation, sequencing, and variant calling

DNA was extracted from dried tissue using Qiagen DNeasy Plant Mini Kits and quantified with a Qiagen QIAxpert microfluidic analyzer (Qiagen Inc., Valencia, CA, USA). We constructed reduced-representation libraries for Illumina sequencing using a ddRADseq method (Parchman et al., 2012; Peterson et al., 2012). The genomic DNA was digested with two restriction enzymes, *EcoRI* and *MseI*, and custom oligos with Illumina base adaptors and unique barcodes were ligated to the digested fragments (ranging from eight to 10 base pairs in length). Ligated fragments were amplified by PCR using a high-fidelity proofreading polymerase (iProof High-Fidelity DNA Polymerase, BioRad Inc., Hercules, CA, USA) and subsequently pooled into a single library. Libraries were size selected for fragments between 350 and 450 bp in length with the Pippin Prep System (Sage Sciences, Beverly, MA) at the University of Texas Genome Sequencing and Analysis Facility (UTGSAF). Sequence data were generated for the full set of individuals using a partial lane of sequencing on the Illumina NovaSeq platform at the UTGSAF.

We used the tapioca pipeline (https://github.com/ncgr/tapioca) and a known contaminant sequence database to identify and discard Illumina primer/adapter sequences and potential biological sequence contaminants (e.g., PhiX, *E. coli*). We then demultiplexed the reads using a custom Perl script that corrects one or two base sequencing errors in barcoded regions, parse reads according to their associated barcode sequence, and trims restriction site-associated bases. Trimmed fastq files for each individual are available at SRA (https:).

Filtered reads were clustered for variant identification and filtering with the software `Stacks v 1.46` (Catchen et al., 2013). We followed a *de novo* assembly approach, using the “denovo_map.pl” module, which allows genotype inference by identifying SNP loci without a reference genome. The parameters were set as follows: the minimum number of identical reads required to call an allele was set to 3 (m = 3), the maximum number of mismatches between loci for individuals was set to 2 (M =2), and the maximum number of mismatches among loci when comparing across individuals was set to 2 (n = 2). These parameters were selected through an optimization process following recommendations from Mastretta-Yanes et al., (2015) and Paris et al., (2017). In brief, we set the optimal m among values ranging 2 to 7 (for M and n = 2) and the optimal M value among values ranging from 2 to 6 for the m optimal value (for n = M). Then, we used the “populations” module in `Stacks` (Catchen et al., 2011, 2013; Rochette et al., 2019) to extract loci that were present in at least 80% of the individuals (--r = 0.80) and with a maximum observed heterozygosity of 0.65 (--max_obs_het = 0.65) and to generate and export the SNP data in vcf format for further analyses. We used `vcftools v 4.2` (Danecek et al., 2011) to estimate the allele frequency, the mean depth per individual, the mean depth per site, and the proportion of missing data per site of the vcf outputs. We explored these statistics in R in order to decide the optimal m, M, and n parameters. Additionally, we filtered the obtained pool of loci using `vcftools v 4.2`. We allowed a maximum missing data of 20 % (--max-missing 0.8), a minimum minor allele frequency of 0.03 (--maf 0.03) and specified a thin value of 5 (--thin 5), which allows that no two sites are within the specified distance from one another. Also, we only included sites with quality scores above 10 (--minQ 10).

### Patterns of population genetic diversity and differentiation

Genetic diversity estimates were calculated in R using the package `hierfstat v 0.5-7` (Goudet, 2005). We used the “basic.stats” function to estimate mean observed heterozygosity (H_o_), mean expected heterozygosity (H_e_), and individual inbreeding coefficients (F_IS_) within each population. Pairwise Nei’s *F*_ST_ (Nei, 1987) and pairwise genetic distances (Nei’s D) were estimated for all pairs of populations with the “genet.dist” function. The confidence intervals over F_IS_ and *F*_ST_ values were estimated using 1000 bootstraps with the “boot.ppfis” function. Additionally, we estimated nucleotide diversity (**π**) and Tajima’s D for each population, using `vcftools v 4.2`.

We further quantified genetic structure within and among populations with an analysis of molecular variance (AMOVA) using the R package `poppr v 2.9.2` (Kamvar et al., 2014). We tested whether most genetic variance was observed among individuals within populations (i.e., no population structure) or between populations (i.e., population structure). The significance of AMOVA results was tested with the function “randtest” from the R package `adegenet v 1.3-1` (Jombart, 2008) using 999 simulations. A linkage disequilibrium (LD) analysis was conducted based on the index of association (I_A_; Brown et al., 1980) and the standardized index of association (r^—^_d_) over all the loci to infer the mode of reproduction within populations. Linkage disequilibrium is expected to be more pronounced in populations engaging in selfing or asexual reproduction in comparison to those mainly reproducing sexually. We used the package `poppr v 2.9.2` (Kamvar et al., 2014) and performed the analysis using 999 permutations. To distinguish between selfing and asexual reproduction as processes leading to low genetic diversity, we estimated relatedness among individuals using `vcftools v 4.2` based on the Manichaikul et al., (2010) approach (-- relatedness2). This method gives information about the relationship of any pair of individuals by assessing their kinship coefficient, which ranges from 0 (no relationship) to 0.50 (self). Individuals were plotted against one another using the “heatmap” function from the R package `stats v 3.3.1` (Team, R. C., 2013).

### Spatial genetic structure

We tested whether populations exhibited isolation by distance (IBD; Wright, 1943) and/or isolation by environment (IBE; Wang & Bradburd, 2014), comparing the pairwise matrices of genetic distances (Nei’s D; see above) with geographic and environmental distances (see environmental data details below) through Mantel tests. We used the “mantel” function from the R package `vegan v 2.5-7` (Oksanen et al., 2013), with Spearman correlation and 9999 permutations. We estimated the geographic distances among populations as haversine distances using the “distm” function of the `geosphere v 1.5-14` R package (Hijmans et al., 2017).

Spatial genetic variation was further assessed using model-free and model-based inference of genetic variation among individuals. First, we inferred population structure and individual ancestries without *a priori* information on sample origin using `ADMIXTURE v 1.3.0` (Alexander and Lange, 2011). We used `PLINK v 1.07` (Purcell et al., 2007) to convert the vcf file into unlinked SNPs (i.e., LD-pruned SNPs) and then ran `ADMIXTURE` with *K* values ranging from 2 to 10. The optimal value of *K* was estimated by evaluating cross-validation errors. Patterns of genetic variation were further summarized by principal component analyses (PCA; Patterson et al., 2006) using the “prcomp” function from the R package `stats v 3.3.1` (Team, R. C., 2013). For each ancestral population (*k*) we indicated the corresponding seed zone of Johnson et al. (2017). We also performed uniform manifold approximation and projection analyses (UMAP; Leland et al., 2018; McInnes et al., 2018) using the “umap” function from the R package `umap v 0.2.7.0` (Konopka, 2020). UMAP has recently been shown to excel at detecting and conveying fine-scale spatial genetic structure of populations (Diaz-Papkovich et al., 2019; Diaz-Papkovich et al., 2021). We ran UMAP with the minimum distance between points in low-dimensional space (MD) ranging from 0.1 to 0.99 and the number of approximate nearest neighbors used to construct the initial high-dimensional graph (NN) ranging from two to 16. Based on an assessment of clustering results across this range of parameter space (Suppl. Figures 2 a-h), we present results here with min_dist = 0.25 and n_neighbors = 16.

We conducted two analyses to visualize variation in differentiation and effective migration across the sampled populations. Both analyses essentially quantify the extent to which differentiation among populations departs from the expectation of isolation by distance. First, we visualized effective migration rates using EEMS (Estimated Effective Migration Surfaces; Petkova et al., 2016). This analysis assigns individuals to the nearest deme, and by using a stepping-stone model, estimates effective migration rates between demes. A genetic dissimilarity matrix was calculated using the bundled bed2diffs script (Petkova et al., 2016). The habitat polygon was obtained manually to include the sampling localities of all the populations, using Google Maps API v 3 Tool (http://www.birdtheme.org/useful/v3tool.html). We chose a deme size of 300 (nDemes parameter) and performed three independent analyses using runeems_snps script, with a burn-in of 100,000,000 (numBurnIter parameter), MCMC length of 200,000,000 (numMCMCIter parameter), and the number of iterations to thin between two writing steps of 999,999 (numThinIter parameter). We combined the results of the three independent runs and plotted the results corresponding to the surfaces of effective diversity (q) and effective migration rates (m) using the R package `rEEMSplots v 0.0.1` (Petkova et al., 2016).

As an additional method to visualize differentiation and effective migration across populations, we employed unbundled principal components (unPC) as a complementary method to EEMS, to reveal potential long-distance migration using the `unPC v 0.1.0` R package (House and Hahn, 2018). UnPC uses principal components in combination with geographic coordinates of samples to create visualizations of genetic differentiation across the landscape. It first calculates the Euclidean distance between PCA coordinates for each pair of populations and then estimates the pairwise geographic distance between populations. The ratio of the genetic distance to the geographic distance for each pair of populations is the unPC value for each pair.

Finally, we analyzed and visualized population structure using a phylogenomic approach. First, we converted the vcf file to fasta format using `vcf2phylip v 2.0` (Ortiz, 2019). We trimmed the fasta alignment to exclude unreliably aligned positions with `trimAl v 1.2` using the “gappyout” method (Capella-Gutiérrez et al., 2009). Then, we ran `IQ-TREE v 1.6.10` (Nguyen et al., 2015) using the “Model Finder Plus” parameter (-m MFP) to determine the best substitution model (choosing the model that minimizes the BIC score), the ascertainment bias correction method (ASC; Lewis, 2001), and the ultrafast bootstrap option with 1000 bootstrap replicates (-bb 1000). We visualized the obtained tree in `Figtree v 1.4.4` (Rambaut, 2018), and indicated the appropriate seed zone for each sampled individual. We then plotted the tree linked with the population’s geographical coordinates after simplifying the tree to retain one sample tip per population using the drop.tip function in the R `APE package v 5.5` (Paradis et al., 2004). We used the “phylo.to.map” function in the R package `phytools v 0.7-80` to plot a map (Revell, 2014), using the dropped tree and the geographical coordinates and choosing the “state” database, again indicating seed zone for each population. To quantify the extent to which seed zones reflect evolutionary history, we tested for phylogenetic signal by estimating Pagel’s λ (Pagel, 1999) for the distribution of seed zones across sampled populations. We used the “fitDiscerte” function from the R package `geiger v 2.0.7` (Harmon et al., 2015). To test the significance of our results, we estimated the log-likelihood if λ = 0 and if λ = 1 using the function “rescale” and did a likelihood ratio test.

### Influence of environmental variation on spatial genetic variation

We conducted genetic-environment association (GEA) analyses using partial redundancy analysis (pRDA) to identify environmental variables that covary with genetic differentiation among populations. Climate environmental variables for each site were obtained from the PRISM database (https://prism.nacse.org) using the “get_prism_normals” function from the R prims library (Hart et al., 2015), with a data range from 1981 to 2010 with an 800 m x 800 m resolution. Following Faske et al., (2021), we converted monthly normals to estimates of potential evapotranspiration, actual evapotranspiration, soil water balance, and climatic water deficit, which have been shown to effectively predict aspects of spatial and distributional variation across plant communities (Barga et al., 2018). Elevation data was acquired from the R library `elevatr v 0.2.0` (Hollister and Shah, 2017). We also included several climatic variables that predicted genecological variation and were used for seed zone delineation in a previous study (Johnson et al. 2017; see Suppl. Tables 2a, b for details). Before any analyses, we examined multicollinearity among the pool of variables using the “pairs.panels” function from the `pyshc v 2.1.9` R package (Revelle, 2015), based on Pearson’s |r| ≤ 0.60, to select the most orthogonal subset of variables possible. We removed all the environmental variables that were highly correlated with other variables, thus reducing the data set from 46 to 8 environmental variables (Table 2). Then, we applied partial redundancy analysis variance partitioning to decompose the contribution of climate, population structure, and geography in explaining genetic variation. We used three sets of variables: 1) climate environmental variables (Table 2); 2) two proxies of genetic structure (population scores along the two previously estimated PC axes); and 3) each population’s coordinates (longitude and latitude). As a response variable, we used the individual-based genotypes (coded as the count of one allele, i.e., 0/1/2). We used the “rda” function from the `vegan v 2.5-7` R package (Oksanen et al., 2013) for pRDA. Following Capblancq and Forester (2021), we first tested the significance of the full RDA model (with all the variables included). Subsequently, explanatory variables were added one by one, using the “ordiR2step” function of the `vegan v 2.5-7` R package, with the following stopping criteria: variable significance of *p* < 0.01, 1000 permutations, and the adjusted R2 of the global model. Then, we performed three different pRDA models: first, a model accounting for environmental variables only (conditioning the model by geography and population structure); second, a model accounting for population structure (conditioning the model by geography and environmental variables); and third, a model accounting for geography (conditioning the model by population structure and environmental variables). We then compared the amount of variance explained by each pRDA to the variance of the full model (including all explanatory variables) to estimate the independent contribution of each set of variables together with any confounding effects induced by collinearity.

**Table 2.**
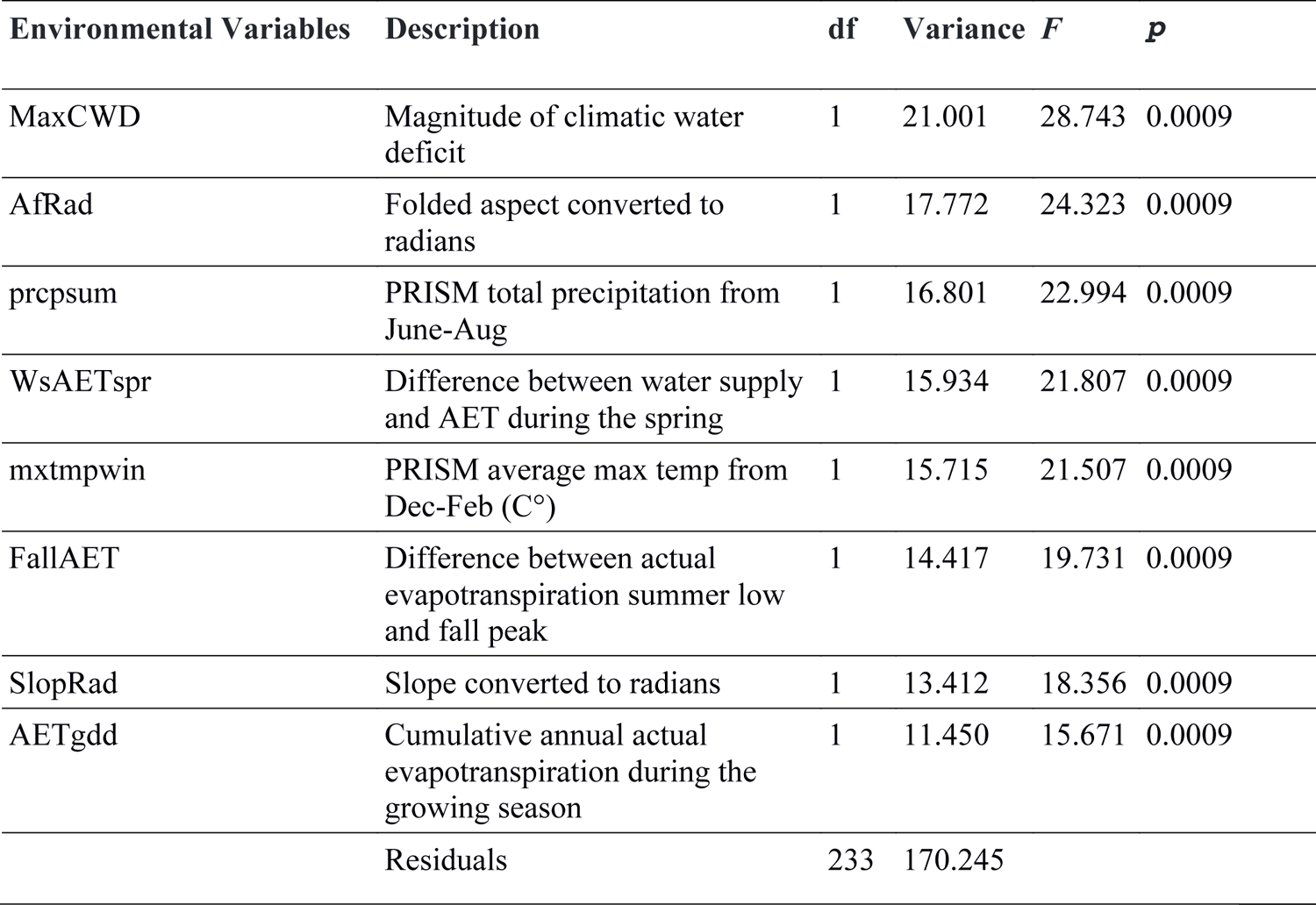
ANOVA results showing the variance explained by each environmental variable in the partial RDA, the *F* value, and the associated *p* value.

| Environmental Variables | Description | df | Variance | $F$ | $P$ |
| --- | --- | --- | --- | --- | --- |
| MaxCWD | Magnitude of climatic water deficit | 1 | 21.001 | 28.743 | 0.0009 |
| AfRad | Folded aspect converted to radians | 1 | 17.772 | 24.323 | 0.0009 |
| prcpsum | PRISM total precipitation from June-Aug | 1 | 16.801 | 22.994 | 0.0009 |
| WsAETspr | Difference between water supply and AET during the spring | 1 | 15.934 | 21.807 | 0.0009 |
| mxtmpwin | PRISM average max temp from Dec-Feb (C°) | 1 | 15.715 | 21.507 | 0.0009 |
| FallAET | Difference between actual evapotranspiration summer low and fall peak | 1 | 14.417 | 19.731 | 0.0009 |
| SlopRad | Slope converted to radians | 1 | 13.412 | 18.356 | 0.0009 |
| AETgdd | Cumulative annual actual evapotranspiration during the growing season | 1 | 11.450 | 15.671 | 0.0009 |
| Residuals |  | 233 | 170.245 |  |  |

Given the results from the above analyses, we conducted pRDA on the genotypic and environmental data to infer the influence of specific environmental variables on spatial genetic structure and detect the genetic signal of local adaptation and its environmental causes. We used a partial RDA (pRDA) conditioning by population structure (PC1 and PC2) and geography (latitude and longitude) to assess whether the degree of adaptive genetic variation among individuals is explained by a particular set of environmental variables. The significance of models and RDA axes, and the proportion of variation explained by each environmental variable were tested with an analysis of variance (ANOVA) and permutation (n = 999), using the “anova.cca” function of the vegan v 2.5-7 R package. Also, we used RDA to identify outlier loci potentially under selection using loadings of SNPs from the first three constrained ordination axes. We used stringent outliers filtering of 3.5 standard deviations (*p* < 0.0005) (Forester et al., 2018). Then, we checked for duplicate candidate loci associated with more than one RDA axis and used Pearson’s correlation (r) to identify the strongest predictor.

## Results

### Patterns of genetic diversity and differentiation

We identified a total of 5,677 SNPs in a subset of 246 individuals that were retained after filtering with mean coverage depth per individual of 21.83. Population genetic statistics were obtained for each population (Table 1). Observed heterozygosity (H_o_) values were lower than the expected (H_e_) in all the cases, indicating low heterozygosity in all the populations studied. As follows, inbreeding coefficient (F_IS_) values were positive for all populations, indicating reduced diversity that is consistent with self-pollination. F_IS_ values were variable across populations but were significantly positive for all populations except DH, EW, SC, SS, and VM, based on bootstrap coefficient intervals. Nucleotide diversity (**π**) estimates were low, congruent with previous results. Moreover, all populations had positive values of Tajima’s D consistent with population size contraction. The index of association (I_A_) and the standardized index of association (r^—^_d_) were different from zero and significant in all cases (*p* < 0.001), indicating elevated levels of linkage disequilibrium consistent with selfing influencing variation within populations. Lastly, 98.58 % of the pairwise combinations among individuals had mean relatedness coefficients of 0, and 1% had a mean relatedness coefficient of 0.33, ranging from 0.005 to 0.47 (considering the 1 % left from comparisons among same individuals) indicating that populations are diverse and do not consist of apomicts (see Suppl. Table 3 and Suppl. Figure 1).

### Spatial genetic structure

AMOVA analyses indicated that 79.66 % (*p* < 0.01) of the observed genetic variance was explained by variation between populations, consistent with strong population differentiation, with the remaining 20.33 % (*p* < 0.01) reflecting variation among individuals within populations (Table 3). The results of the Mantel test indicated a positive association between geographic distance and genetic distance (IBD: Mantel statistic *r*: 0.257, *p*: 0.026; Figure 2a) and between environmental and geographic distances (Mantel *r*: 0.238, *p*: 0.025; Figure 2c). No significant association was found between environmental and genetic distances (Mantel statistic *r*: −0.001, *p*: 0.437; Figure 2b).

**Table 3.**
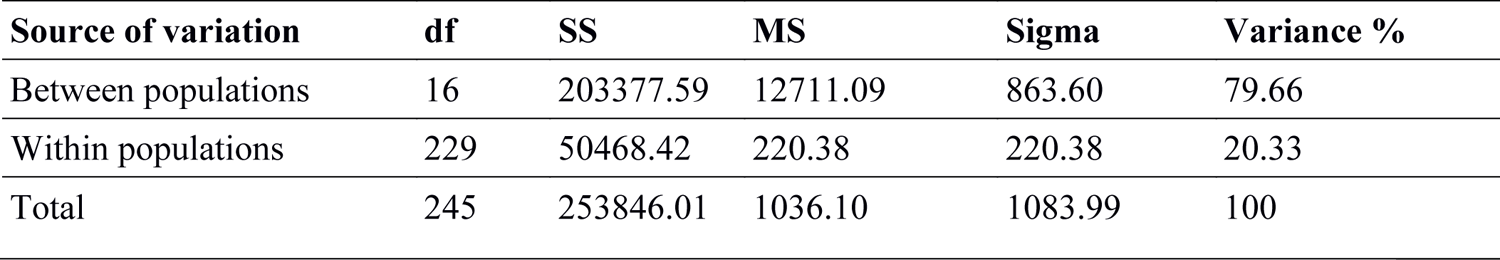
Molecular analysis of variance (AMOVA) results for the 17 populations of *Achnatherum thurberianum*. Here, we provide the degrees of freedom (df), sum of squares (SS), mean squares (MS), total variance (sigma), and the percentage of variance explained by each source of variation (Variance %).

| Source of variation | df | SS | MS | Sigma | Variance % |
| --- | --- | --- | --- | --- | --- |
| Between populations | 16 | 203377.59 | 12711.09 | 863.60 | 79.66 |
| Within populations | 229 | 50468.42 | 220.38 | 220.38 | 20.33 |
| Total | 245 | 253846.01 | 1036.10 | 1083.99 | 100 |

Population pairwise *F*_ST_ values were significant in all pairwise comparisons (mean: 0.18, range: 0.02 – 0.31), even for those involving populations that were highly spatially proximate, indicating significant genetic differentiation between populations (Suppl. Table 4). PCA revealed three strongly separated population genetic clusters (Figure 1b), but also suggested a high degree of identifiability of individuals from most populations. The first two principal components accounted for 16.75 and 13.27% of the variation. One cluster grouped the eastern populations (AH, AS, BM, FR, SC) and the HO population, the second cluster grouped the western populations (BV, GB, JC, LV, MD, PL, PT, SS, and VM), and lastly, the third cluster grouped EW and DH. The UMAP analyses revealed a striking fine-scale population genetic structure in which all individuals for each population clustered tightly together (note that the distances among UMAP clusters do not represent genetic differentiation among them; Figure 1c). UMAP analyses across the ranges of the minimum distance (MD) and the number of nearest neighbors (NN) parameters also produced generally consistent clustering patterns (Suppl. Figures 2a-h).

`ADMIXTURE` identified *K* = 8 as the most likely number of clusters among the 17 populations sampled (Figure 1d), based on cross-validation error values (Suppl. Figure 3). Ancestry coefficient estimates for this analysis produced a similar pattern to the PCA and the UMAP results. Eastern populations formed an ancestry cluster (AH, AS, BM, FR, and SC), while western populations split into two different ancestry clusters (GB, JC, MD, and SS, in one, and PL, PT, and VM, in another). Lastly, the BV, DH, EW, HO, and LV populations were assigned to additional single clusters, reflecting relatively stronger differentiation of these populations in relation to those within the larger ancestral clusters above. Notably, populations from individual seed zones of Johnson et al. (2017) were commonly assigned to multiple ancestral groups in the `ADMIXTURE` results (Figure 1d, e.g., populations in seed zones represented by circles belong to three different ancestry groups), illustrating discordance among evolutionary history and seed zones delineated with phenotype-environment associations. The two other most likely *K* values, *K* = 9 and *K* = 10, generated similar patterns of cluster membership and similar discordance among seed zones and ancestry (see Suppl. Figure 3b).

The EEMS results suggested effective migration patterns congruent with previous results. Some populations were connected by higher migration rates (m) than expected under isolation by distance. For example, HO and AH, while separated by approximately 350 km, were connected with high effective migration (Figure 1, Figures 3b, c). Moreover, other groups of populations seem to have resistance barriers to gene flow despite being highly proximate geographically (Figure 3b). For example, PL and PT appear to be distinguished by low effective migration rates despite being separated by only 19 km. Results of unPC analyses (Suppl. Table 5) were broadly consistent with those from the EEMS, suggesting the same regions of low and high effective migration (Figure 3c).

**Figure 3.**
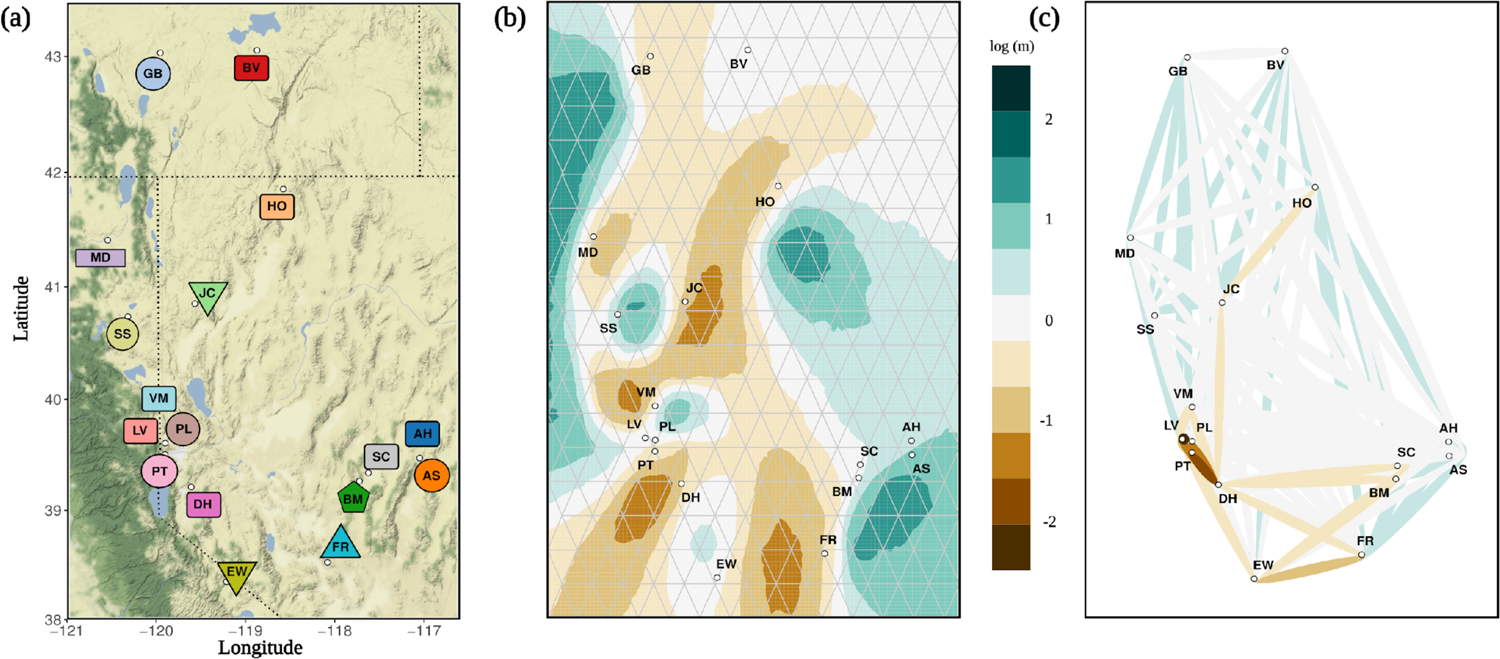
Landscape genetic differentiation for the 17 *Achnatherum thurberianum* populations. a) Map of the sampled locations with each population (colors) and seed zone (symbols) code (Table 1). b) Estimated effective migration surface (EEMS) plot depicting estimated migration rates (in log 10 scale) that deviate from isolation by distance expectations. Brown represents areas of low effective migration relative to the average, and green represents areas of higher effective migration. c) Unbundled principal components (unPC) representation; higher unPC values, representing greater differentiation than expected among populations, are colored in progressively darker shades of brown, while lower unPC values, representing lower differentiation among populations, are colored in progressively darker shades of green. The unPC values near the mean of the distribution are colored in white.

The maximum likelihood tree in which all main clades yielded branch support higher than 99 produced topologies similarly illustrated pronounced population divergence. Similar to the patterns of clustering in PCA and `ADMIXTURE` analyses above, tree topology resolved four main clades; the eastern populations (AH, AS, BM, FR, and SC) with HO, the BV population, the EW and DH populations, and lastly, the western populations (GB, JC, MD, SS, LV, PL, PT, and VM) (Figures 4a, b). Similar to evidence for population identifiability in UMAP analyses, individuals from the same populations predominantly clustered together in the maximum likelihood tree, further illustrating population differentiation at fine spatial scales. Consistent with ancestry based analyses above, there was no evidence for phylogenetic signal for seed zones (λ = 0.000); populations from the same seed zones often appeared in multiple distantly related clades (e.g., DH and EW, or PL and VM, Figures 4 a, b).

**Figure 4.**
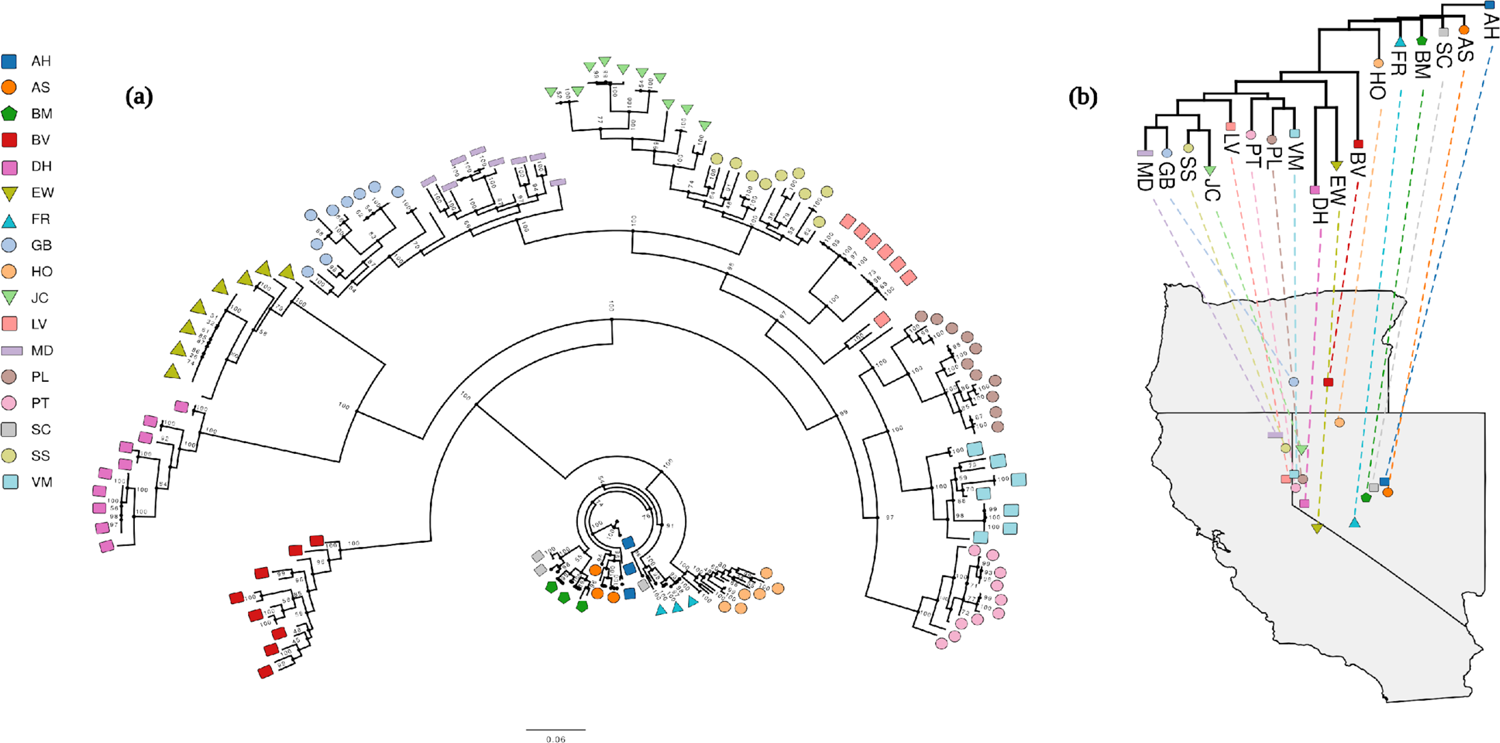
Population phylogenetic differentiation of the 17 *Achnatherum thurberianum* populations. a) Maximum likelihood topology inferred with `IQ-TREE`. The scale bar represents the expected number of nucleotide substitutions per site. Ultrafast bootstrap support values (UFBS) are indicated at the nodes. b) Projected phylogeny onto the geographic map showing each population’s location. One individual is represented for each population to minimize overlap. The different symbols in both panels correspond to the different seed zones studied (Table 1).

### Genetic-environment association analyses

The pool of environmental variables was reduced from 46 to 8 after removing highly correlated variables (based on Pearson’s |*r*| ≤ 0.60; note that the environmental variables used for seed zone delineation in Johnson et al. (2017), were not included in the analyses to control for multicollinearity, but some of them were correlated with the eight variables selected from our analyses see Suppl. Table 2c). Results from the pRDA provided evidence that specific environmental variables may influence spatial patterns of genetic variation. In particular, the climatic variables explained 23% of the total genetic variation (39% of the variance explained by the full model), suggesting an association between genetic variation and environmental gradients (IBE) (Table 4). The environmental variables with the greatest explained variance in the pRDA were the magnitude of climate water deficit (MaxCWD; 21.01 %, *p* < 0.001) and the folded aspect (AfRad; 17.80 %, *p* < 0.001) (Table 2; Figure 5). The first two constrained axes were significant (*p* < 0.001), explaining 24.63 % and 23.44 % of the total variation (Figure 5). We identified ten loci across the environmental variables associated with the second and third RDA axes (Suppl. Table 2d). Four of these ten loci were associated with the difference between water supply and actual evapotranspiration during the spring (WsAETspr), two with the maximum temperature during the winter season (mxtmpwin), and the four remaining were associated with the cumulative annual actual evapotranspiration during the growing season (AETgdd), the difference between actual evapotranspiration summer low and fall peak (FallAET), the magnitude of climatic water deficit (MaxCWD), and the slope (SlopRad). One of the variables strongly associated with genetic variation the maximum temperature during the winter season (mxtmpwin; Table 2) was highly correlated with the mean average temperature (MAT; Pearson’s |*r*| = 0.82), which was among the variables most strongly predicting genecological variation in Johnson et al. (2017).

**Figure 5.**
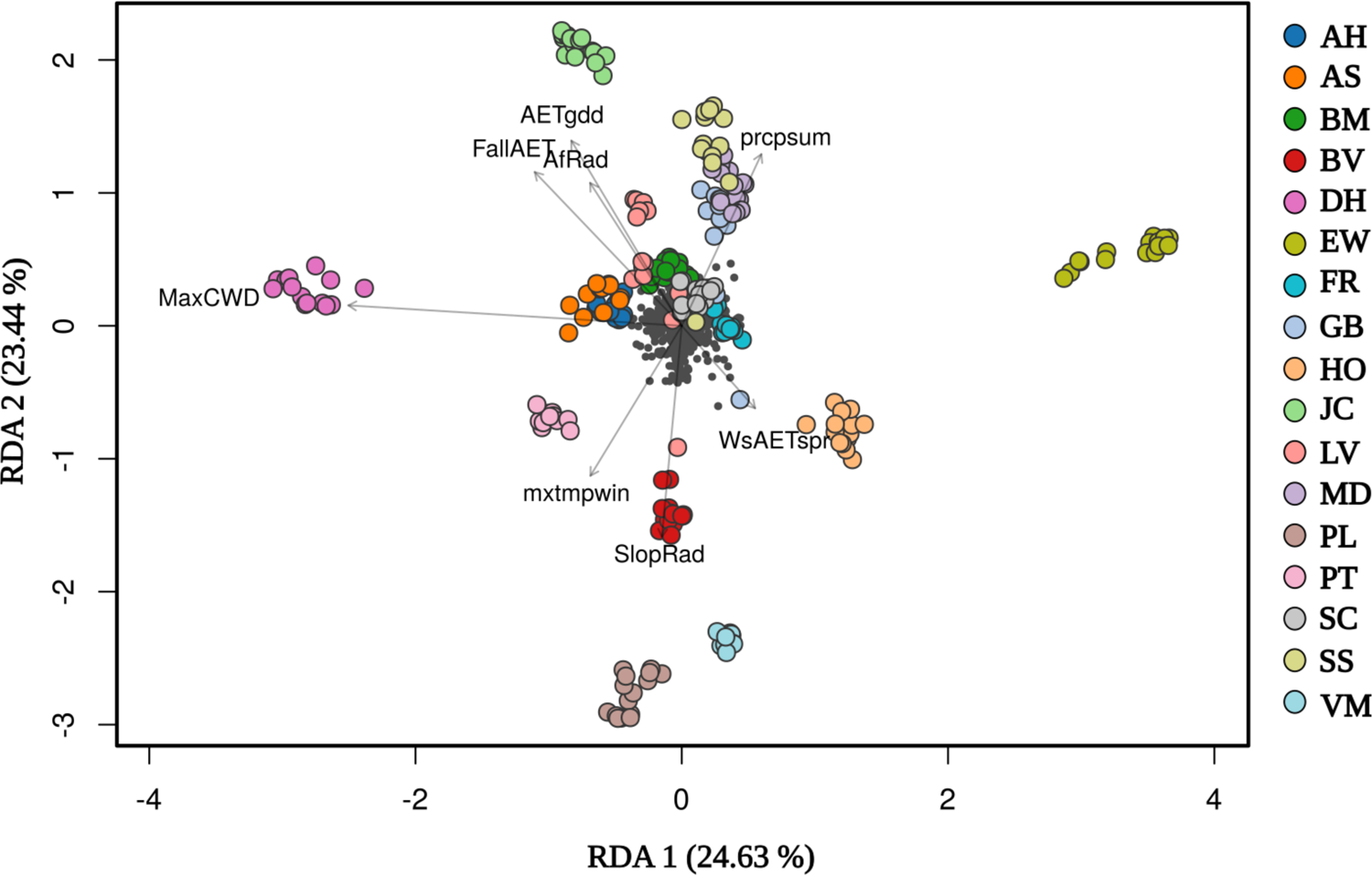
Redundancy analysis (RDA) plot depicting the environmental variables studied significantly associated with genetic variation. The direction and length of arrows correspond to the loadings of each variable on the two RDA axes.

**Table 4.**
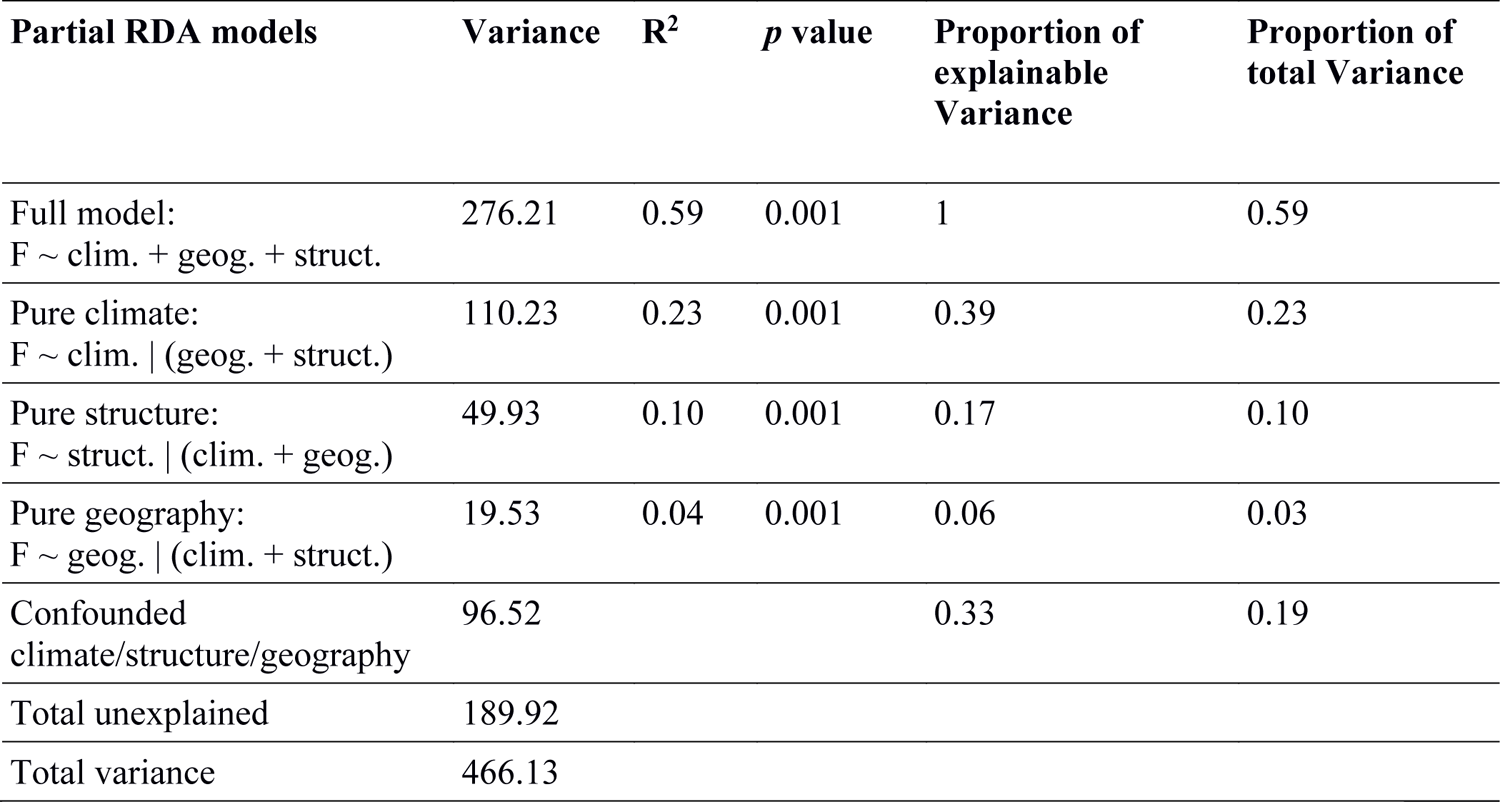
The influence of climate, structure, and geography on genetic variation decomposed with partial RDA. The proportion of explainable variance represents the total constrained variation explained by the full model.

## Discussion

Understanding the nature of genetic variation in native plants is crucial not only for understanding the origin and maintenance of diversity but also for conserving and restoring populations. Our analyses of population genetic variation in Thurber’s needlegrass (*Achnatherum thurberianum*), a widespread bunchgrass in the Great Basin, illustrated strong regional differentiation as well as remarkably fine-scale spatial genetic structure among populations. These patterns, along with low levels of genetic diversity within populations, high inbreeding coefficients, and elevated linkage disequilibrium, are consistent with self-pollination reducing genetic diversity and contributing to spatial genetic structure. Despite low genetic diversity within the sampled populations, our analyses indicated that environmental variation has shaped spatial genetic structure and influenced local adaptation across *A. thurberianum* populations. Our results suggested the potential for local adaptation driven particularly by climate water deficit, aspect, and summer precipitation. Notably, our results illustrated previously unidentified differences in evolutionary history within seed zones proposed for *A. thurberianum* based on phenotype-environment associations (Johnson et al. 2017). Altogether, our results suggest that numerous factors have shaped the spatial genetic structure of *A. thurberianum* across fine geographic and environmental scales and provide baseline information that may be of value for restoration, including allowing managers to consider both phenotypic variation and evolutionary history when making seed source decisions.

Strongly congruent results across multiple analyses (*F*_ST_, PCA, `ADMIXTURE`, phylogenomic analyses, AMOVA, and UMAP) illustrated spatial genomic structure across both broad geographic regions and among proximate populations at finer scales than might otherwise be expected for a primarily wind-dispersed species (Linder et al., 2018). For example, at broad regional scales, our analyses consistently illustrated pronounced differentiation among eastern and western groups of populations (Figures 1, 4) including differentiation of two populations at the southwestern limit of sampling. Interestingly, the HO population clustered within the eastern populations, despite being geographically distant and in closer proximity to populations in the northwestern portion of the sampled area (e.g., JC; see Figures 1a, b). These broad patterns of regional differentiation could be explained by historical habitat contractions during the multiple Pleistocene glacial cycles that shaped the Great Basin landscape (Beck and Jones, 1997). In particular, during the Last Glacial Maximum and throughout the last deglaciation, the Great Basin region was marked by the formation of multiple extensive lacustrine systems (Tchakerian and Lancaster, 2002; Lyle et al., 2012), which would have reduced the availability of suitable habitat for *A. thurberianum*. More specifically, a large portion of our study area was historically occupied by Lake Lahontan (38.75– 40.75°N, 117.5–120.5°W; Russell, 1885; Matsubara and Howard, 2009), which may have played a central role in shaping regional patterns of marked differentiation in Thurber’s needlegrass. Consistent with such history, estimates of effective migration (EEMS and unPC) indicated barriers to gene flow among eastern and western groups of populations, which continue to be separated by low-elevation playas that are inhospitable to this plant (Figures 3b, c). Further, positive values of Tajima’s D for each population indicated past population contraction (Table 1; Tajima, 1989). Temperate grass species of Europe (Hensen et al., 2010; Blanco-Pastor et al., 2019; Blanco-Pastor et al., 2021), Africa (Mairal et al., 2021), and North America (Avendaño-González et al., 2019; Palacio-Mejía et al., 2021) demonstrate comparable regional patterns of differentiation and gene flow also influenced by Pleistocene glacial cycles.

In addition to the larger-scale geographic patterns, genetic differentiation across smaller geographic scales was evident in phylogenetic analyses, where individuals clustered largely by population, and in the UMAP analyses, where individuals formed population-specific clusters (Figure 1c). UMAP analyses provided remarkable resolution of spatial genetic structure, with all *A. thurberianum* individuals having 100% identifiability to their population of origin. While parameter settings can influence the UMAP depiction of clustering (MD and NN; Diaz-Papkovich et al., 2021), our analyses across a range of two parameters produced largely congruent results with minor variation in the strength of clustering (Suppl. Figure 2). These results suggest that the degree of differentiation among populations seen here could possibly be used to retroactively identify the constituents of historically seeded populations with a high degree of certainty. Estimates of effective migration (EEMS and unPC) also appeared strongly reduced among a number of geographically proximate populations (PT and DH separated by 58 km, or EW and FR separated by 127 km, for example, Figures 3b, c). While gene flow among spatially proximate populations can be high in some wind-dispersed grasses (Vogel et al., 2009; Stritt et al., 2022), pronounced spatial genetic structure has also been reported in Eurasian species of Stipeae (Wagner et al., 2011; Durka et al., 2013). Patterns of population differentiation and identifiability across both large and small geographic scales indicates that genetic variation in *A. thurberianum* has been shaped by a combination of historical isolation, local adaptation to environment, as well as life history variation.

Species that rely strongly on self-fertilization cannot maintain high levels of genetic diversity within populations through frequent pollen movement and therefore tend to have low genetic diversity (Hamrick and Godt, 1996; Honnay and Jacquemyn, 2007; Durka et al., 2013; Huang et al., 2021). Moreover, selfing plant species can be prone to developing fine-scale spatial genetic structure due to reduction in effective population sizes and effective migration (Volis et al., 2010; Huang et al., 2021). The populations studied here were characterized by low levels of genetic diversity (mean H_e_: 0.04; mean H_o_: 0.02; mean π: 0.05), strongly positive inbreeding coefficient estimates (F_IS_), and a high degree of linkage disequilibrium (r^—^_d_ and I_A_) (Table 1), all of which are consistent with a mating system dominated by self-fertilization. While studies directly assessing the mating system of *A. thurberianum* are lacking, many closely related species from the Poaceae family have been described as self-fertilized (Jones and Nielson, 1989; Arnesen et al., 2017; Marques et al., 2017; Stritt et at. 2022). Indeed, grass species employing self-fertilization have been commonly documented to have low genetic diversity and pronounced spatial genetic structure (Dell’Acqua et al., 2014; Marques et al., 2017; Guo et al., 2017), presumably due to small effective population sizes and genetic drift in the absence of much gene flow. Our results suggest high rates of self-fertilization in *A. thurberianum* have likely contributed to its pronounced spatial genetic structure in the western Great Basin.

Although mating system and low genetic diversity have allowed genetic drift to influence spatial genetic structure, environmental variation also appears to have contributed to genetic variation in *A. thurberianum*. The environmental variables influencing spatial genetic variation in *A. thurberianum* are known to predict genetic and phenotypic variation across populations of other plant species (Dilts et al., 2015; Barga et al., 2018; Faske et al., 2021) and have been commonly implicated as underlying local adaptation in genecological studies of Great Basin plant species (Johnson et al. 2017; Baughman et al., 2019). Variance partitioning with partial RDA illustrated that neutral genetic structure, geography, and environmental variation together explained a substantial proportion of genetic variance (Table 4). However, the individual contribution of environmental variation was substantially stronger than that of both genetic structure and geography (Table 4). Variation in environmental variables (Table 2), especially climate water deficit, aspect, and summer precipitation, explained a substantial proportion of genetic variation and influenced spatial genetic structure (Table 2, Figure 5). The pronounced environmental heterogeneity of the Great Basin (Figure 2c) could commonly give rise to Isolation by Environment (IBE), where environmental transitions act as barriers to gene flow or where local adaptation to different environments leads to low migrant fitness (Shafer and Wolf, 2013; Wang and Bradburd, 2014). Genomic evidence for environment contributing to spatial genetic structure and perhaps local adaptation is consistent with results from a common garden study of phenotypic variation across 66 Great Basin *A. thurberianum* populations that inferred local adaptation to environment as a basis for delineating seed zones (Johnson et al. 2017).

Given the strong influence of environmental variables on genetic variation observed here and in Johnson et al. (2017), along with the topographically and environmentally heterogeneous nature of this region, we suspected that proposed seed zones might span populations from distantly related clades. Indeed, geographically differentiated groups of populations commonly crossed multiple seed zones, and specific seed zones were represented across distantly related clades (Figures 1, 4). It is worth noting that our sampling focused on an area that was not well represented in the sampling design for the phenotype-based seed zone development (Johnson et al. 2017), which might indicate that caution is warranted when designating seed zones outside areas of extensive sampling. Convergent evolution, or adaptation to parallel environmental conditions across divergent lineages, is well documented across diverse groups of plants (e.g., Rellstab et al. 2020; Xu et al. 2020), and this possibility should be considered for seed zone design in the Great Basin, especially if particular geographic regions show consistent barriers to gene flow for multiple species. Indeed, other plant species from the Great Basin have been found to have similar patterns of population genetic structure in the vicinity of historic Lake Lahontan (e.g., Faske et al., 2021). Convergent evolution could have consequences for restoration if outbreeding depression (reviewed in Edmands, 2007) or genetic incompatibilities (Etterson et al. 2007) occur in seed mixes containing distantly related populations delineated as part of a single seed zone.

In addition to its relevance for seed zones, a general understanding of genetic diversity, population differentiation, and local adaptation in *A. thurberianum* could have utility for guiding ecological restoration (Mijangos et al., 2015; Breed et al., 2019). Populations exhibited very low levels of standing variation, presumably due to high self-fertilization rates, which likely contributed to population differentiation at such fine spatial scales. Low diversity can be a concern when sourcing seeds for restoration, as genetic diversity is often viewed as a proxy for evolutionary potential (Ellstrand and Elam, 1993), and some have proposed mixing populations of low-diversity species as a way to increase the chances of long-term persistence of restored populations (Bischoff et al., 2010; Bucharova et al., 2019; St. Clair et al., 2020). While this may lead to risks of outbreeding depression in some species, this might be reduced for highly-selfing species like *A. thurberianum* (Jones and Nielson, 1989). Despite very low diversity, our analyses indicated that environmental variation has shaped spatial genetic structure and influenced local adaptation across *A. thurberianum* populations of the Great Basin. Generally, consistent results of inference of local adaptation in our study with that from phenotypic analyses of genecological experiments (Johnson et al., 2017; Baughman et al., 2019) highlight the utility of population genomic analyses for characterizing the environmental variables contributing to local adaptation while additionally characterizing levels of diversity and differentiation across space, all of which have value for guiding provenance and seed sourcing (Breed et al., 2019; Rossetto et al., 2019).

## Data Archiving Statement

The trimmed vcf file and scripts used for analyses can be found at the Dryad Digital Repository: https://doi.org/10.5061/dryad.pvmcvdnpn. The raw data from this project were submitted to NCBI Sequence Read Archive (SRA) and can be found by the BioProject ID PRJNA849003. The individual fastq files for each population can be found under the following accession numbers: AH (SRR19646741), AS (SRR19646740), BM (SRR19646732), BV (SRR19646731), DH (SRR19646730), EW (SRR19646729), FR (SRR19646728), GB (SRR19646727), HO (SRR19646726), JC (SRR19646725), LV (SRR19646739), MD (SRR19646738), PL (SRR19646737), PT (SRR19646736), SC (SRR19646735), SS (SRR19646734), VM (SRR19646733).

## Acknowledgments

The authors thank Fred Edwards, Trevor Faske, Joshua Hallas, Katie Uckele, and Joshua P. Jahner for helpful discussion in support of project design and analyses. Owen Baughman, Lana Sheta, Sage Ellis, and Meagan O’Farrell assisted with the collection of plant material. This work was supported by funding from the U.S. Department of Agriculture (Award Number: 2017-67019-26336) and the U.S. Bureau of Land Management (Award Numbers: L16AC00318 and L19AC00013).

