## Supplementary Material for "Fine-scale spatial genetic structure in a locally abundant native bunchgrass (*Achnatherum thurberianum*) including distinct lineages revealed within seed transfer zones"

### List of Supplementary Tables:

**Table 1.** Relevant environmental variables for each population sampled and seed zones information.

**Table 2.** Summary statistics of environmental variables (attached as xls).

**Table 3.** Coefficient of relatedness among individual pairwise comparisons (attached as csv).

**Table 4.** Pairwise Fst values and their bootstrap values (attached as csv).

**Table 5.** UnPC scores (attached as csv).

### List of Supplementary Figures:

**Figure 1.** Heat map showing relatedness among individuals.

**Figure 2.** UMAP plots with the minimum distance between points in low-dimensional space (min_dist = minimum distance parameter) ranging from 0.1 to 0.99 and the number of approximate nearest neighbors used to construct the initial high-dimensional graph (n_neighbors = number of nearest neighbors' parameter) ranging from two to 16.

**Figure 3.** Cross-validation errors values from and ADMIXTURE plots for K 9 and K 10.

**Table 1.** Population's name and US State where the populations were sampled, two letters population abbreviation, and number of individuals sampled for each population (N). Also, relevant environmental variables are shown for each population: elevation (in meters), mean annual temperature, annual maximum temperature, and annual minimum temperature (in centigrade degrees), and annual mean precipitation (in liters).

| Population name, State | Population abbreviation | N | Elevation (m) | Annual mean temperature (C°) | Annual maximum temperature (C°) | Annual minimum temperature (C°) | Annual mean precipitation (l) |
|---|---|---|---|---|---|---|---|
| Austin Hwy, NV | AH | 14 | 1755 | 8.35 | 17.52 | -0.83 | 0.21 |
| Austin Summit, NV | AS | 15 | 2480 | 7.72 | 13.87 | 1.56 | 0.38 |
| Bald Mountain Canyon, NV | BM | 18 | 2245 | 6.85 | 14.64 | -0.95 | 0.31 |
| Buena Vista, OR | BV | 15 | 1277 | 8.02 | 15.98 | 0.05 | 0.26 |
| Dayton Hill, NV | DH | 14 | 1400 | 10.54 | 19.09 | 1.98 | 0.26 |
| East Walker, CA | EW | 15 | 2043 | 8.01 | 16.79 | -0.49 | 0.29 |
| Finger Rock, NV | FR | 14 | 2129 | 8.45 | 15.43 | 1.46 | 0.25 |
| Grey's Butte, OR | GB | 14 | 1586 | 7.51 | 14.98 | 0.03 | 0.25 |
| Hwy 140, NV | HO | 15 | 1423 | 9.01 | 16.69 | 1.33 | 0.24 |
| Jones Canyon, NV | JC | 15 | 1440 | 9.25 | 17.32 | 1.18 | 0.31 |
| Long Valley, CA | LV | 14 | 1660 | 9.02 | 17.02 | 1.02 | 0.34 |
| Modoc, CA | MD | 15 | 1333 | 8.51 | 16.92 | 0.10 | 0.32 |
| Patagonia, NV | PT | 15 | 1421 | 10.72 | 18.97 | 2.46 | 0.29 |
| Peavine Low, NV | PL | 12 | 1724 | 9.09 | 16.76 | 1.41 | 0.35 |
| Smith Creek, NV | SC | 15 | 2050 | 7.73 | 16.20 | -0.74 | 0.25 |
| Spanish Springs, CA | SS | 14 | 1645 | 7.54 | 15.58 | -0.51 | 0.25 |
| Virginia Mountains, NV | VM | 12 | 1503 | 9.64 | 17.56 | 1.71 | 0.33 |

**Figure 1.** Heat map of relatedness among the 246 individuals studied with a dendrogram showing clusterization among related individuals. The degree of relatedness is indicated by colors, ranging from light yellow (no relatedness) to dark red (strong relatedness).

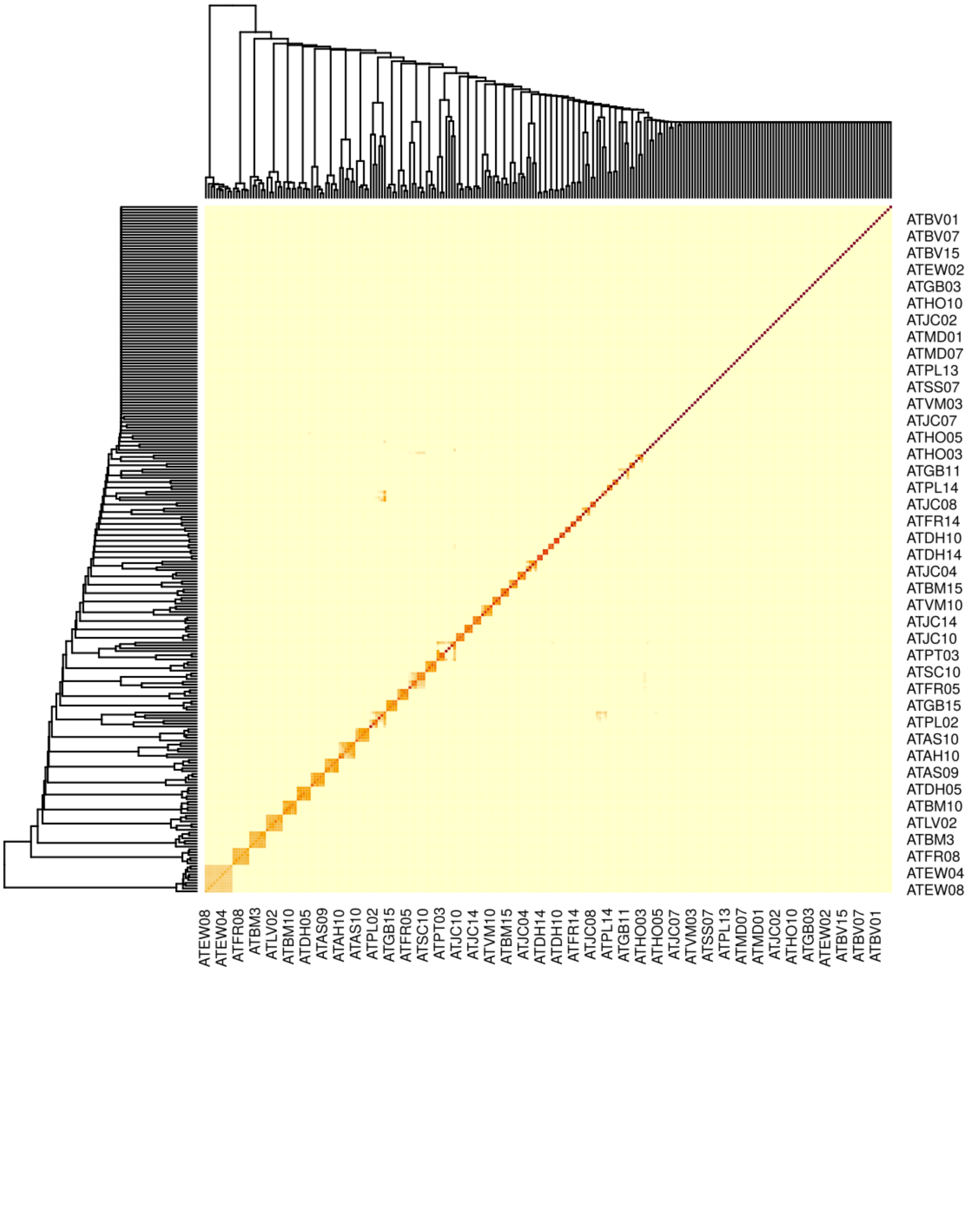

**Figure 2.** UMAP plots for the 17 *Achnatherum thurberianum* populations with the minimum distance between points in low-dimensional space (min_dist = minimum distance parameter) ranging from 0.1 to 0.99 and the number of approximate nearest neighbors used to construct the initial high-dimensional graph (n_neighbors = number of the nearest neighbors parameter) ranging from two to 16. a) UMAP plots with n_neighbors = 2, and min_dist ranging from 0.1 to 0.99. b) UMAP plots with n_neighbors = 4, and min_dist ranging from 0.1 to 0.99. c) UMAP plots with n_neighbors = 6, and min_dist ranging from 0.1 to 0.99. d) UMAP plots with n_neighbors = 8, and min_dist ranging from 0.1 to 0.99. e) UMAP plots with n_neighbors = 10, and min_dist ranging from 0.1 to 0.99. f) UMAP plots with n_neighbors = 12, and min_dist ranging from 0.1 to 0.99. g) UMAP plots with n_neighbors = 14, and min_dist ranging from 0.1 to 0.99. h) UMAP plots with n_neighbors = 16, and min_dist ranging from 0.1 to 0.99.

**a)**

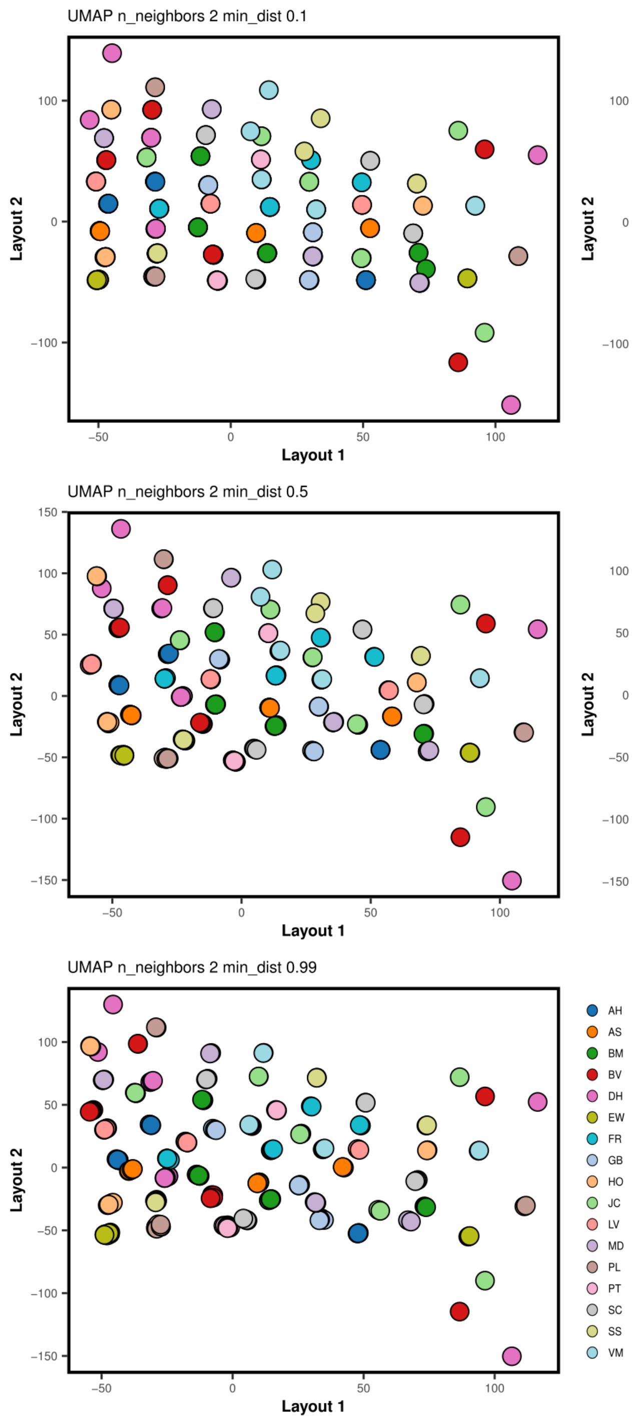
UMAP n_neighbors 2 min_dist 0.1
UMAP n_neighbors 2 min_dist 0.25
UMAP n_neighbors 2 min_dist 0.5
UMAP n_neighbors 2 min_dist 0.8
UMAP n_neighbors 2 min_dist 0.99
Layout 1
Layout 2
−50
0
50
100
150
−100
−150
AH
AS
BM
BV
DH
EW
FR
GB
HO
JC
LV
MD
PL
PT
SC
SS
VM

**b)**

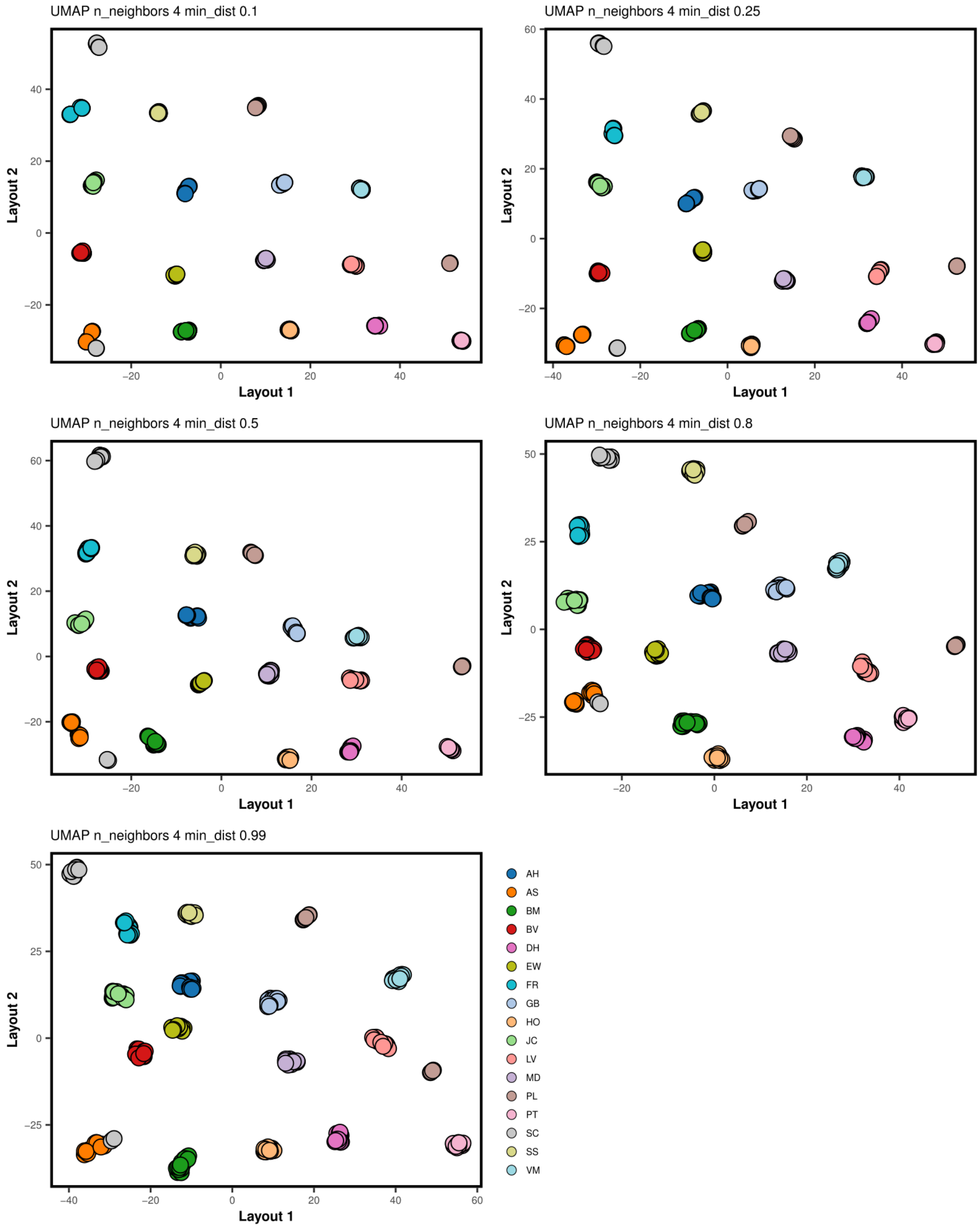
UMAP n_neighbors 4 min_dist 0.1
UMAP n_neighbors 4 min_dist 0.25
UMAP n_neighbors 4 min_dist 0.5
UMAP n_neighbors 4 min_dist 0.8
UMAP n_neighbors 4 min_dist 0.99
Layout 1
Layout 2
AH
AS
BM
BV
DH
EW
FR
GB
HO
JC
LV
MD
PL
PT
SC
SS
VM

**c)**

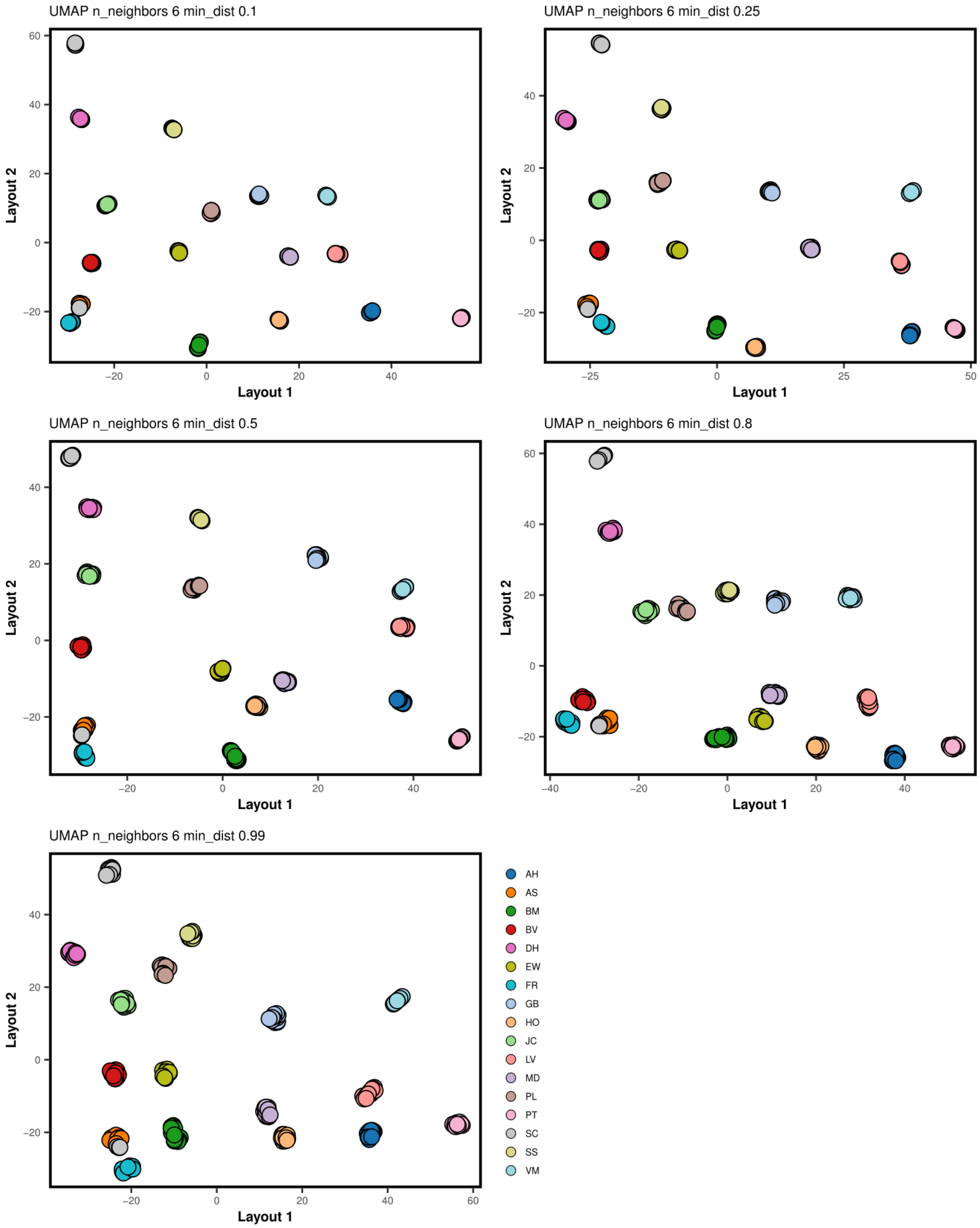
UMAP n_neighbors 6 min_dist 0.1
UMAP n_neighbors 6 min_dist 0.25
UMAP n_neighbors 6 min_dist 0.5
UMAP n_neighbors 6 min_dist 0.8
UMAP n_neighbors 6 min_dist 0.99
Layout 1
Layout 2
AH
AS
BM
BV
DH
EW
FR
GB
HO
JC
LV
MD
PL
PT
SC
SS
VM

d)

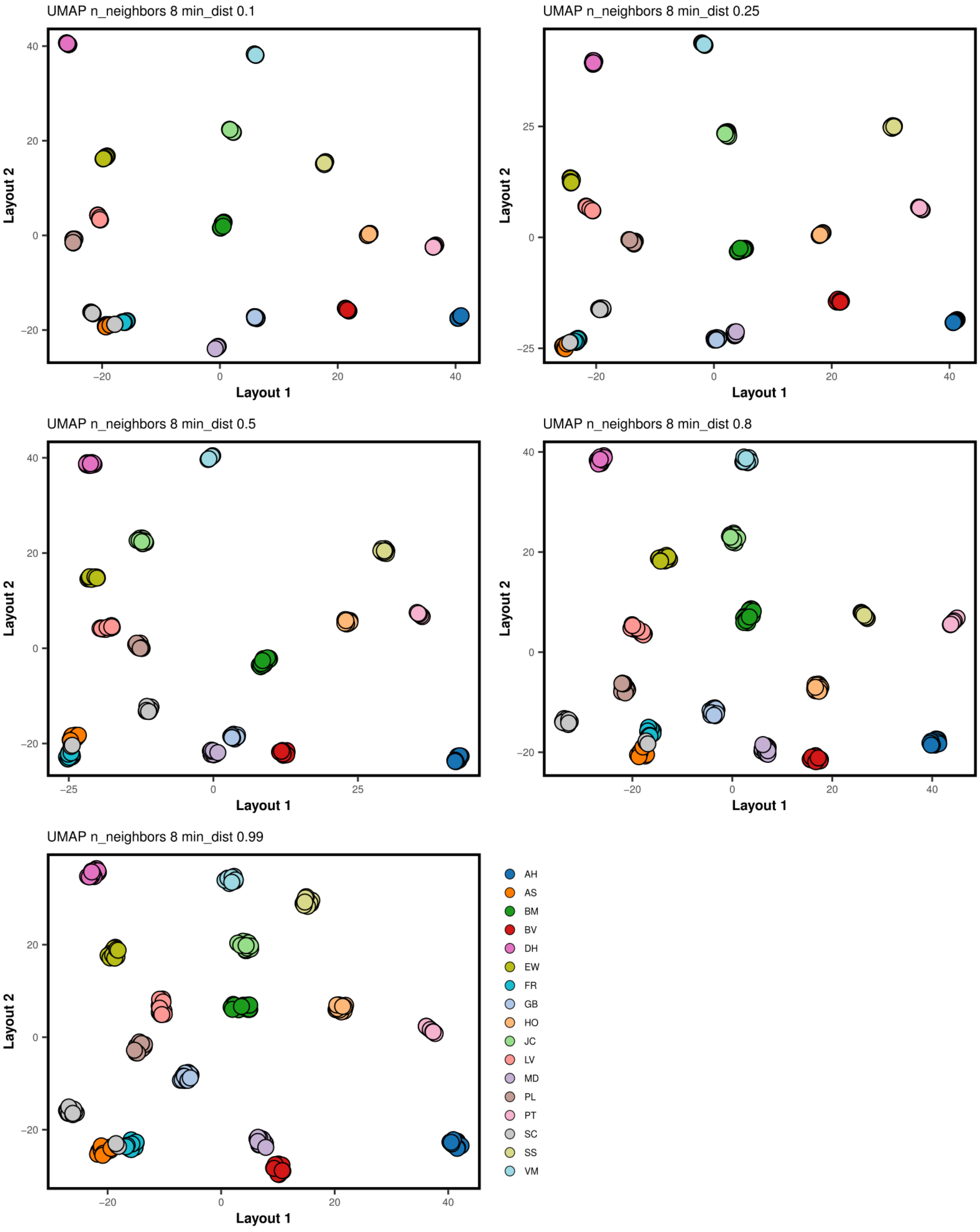

UMAP n_neighbors 8 min_dist 0.1
UMAP n_neighbors 8 min_dist 0.25
UMAP n_neighbors 8 min_dist 0.5
UMAP n_neighbors 8 min_dist 0.8
UMAP n_neighbors 8 min_dist 0.99
Layout 1
Layout 2
AH
AS
BM
BV
DH
EW
FR
GB
HO
JC
LV
MD
PL
PT
SC
SS
VM

**e)**

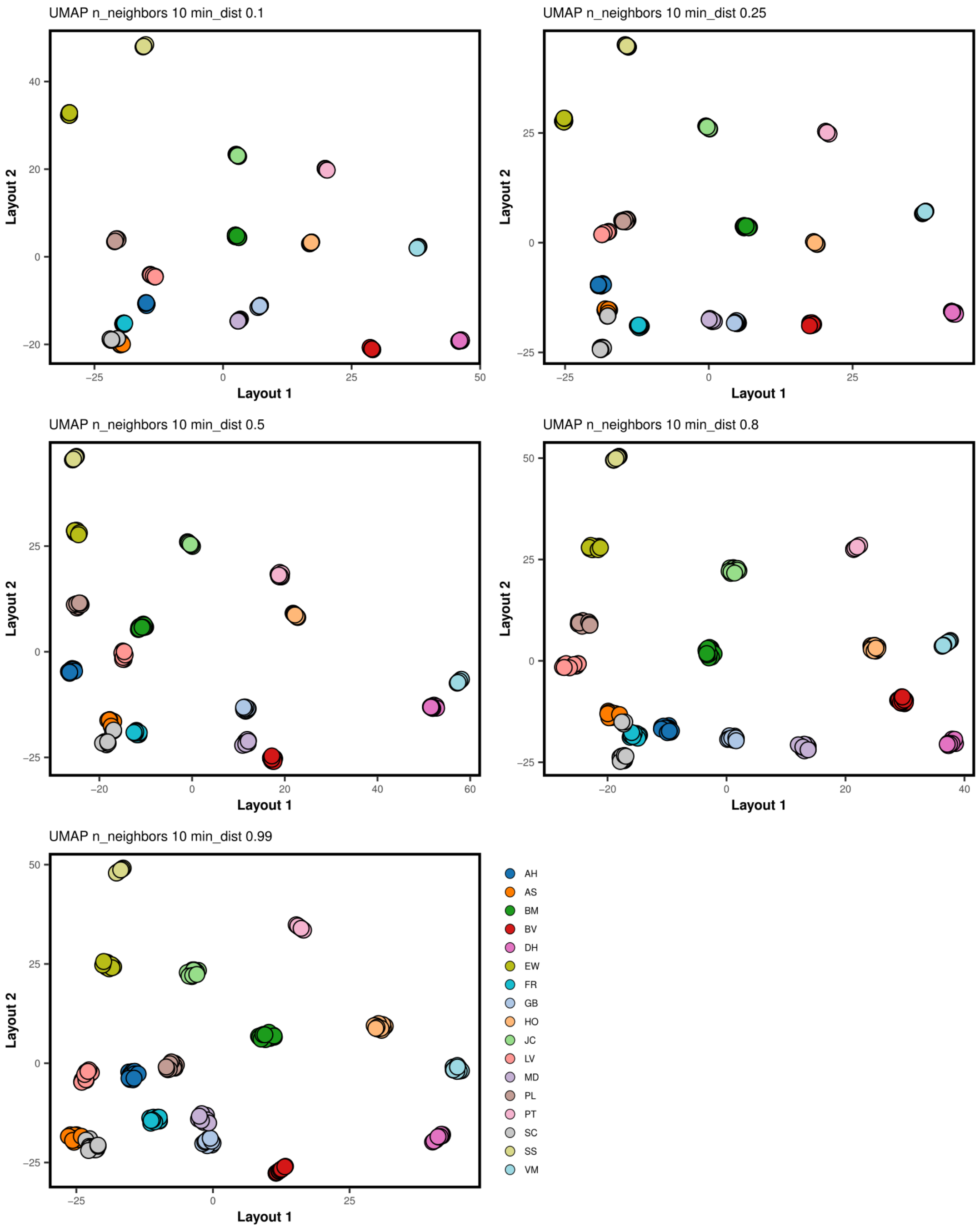

UMAP n_neighbors 10 min_dist 0.1
UMAP n_neighbors 10 min_dist 0.25
UMAP n_neighbors 10 min_dist 0.5
UMAP n_neighbors 10 min_dist 0.8
UMAP n_neighbors 10 min_dist 0.99
Layout 1
Layout 2
AH
AS
BM
BV
DH
EW
FR
GB
HO
JC
LV
MD
PL
PT
SC
SS
VM

**f)**

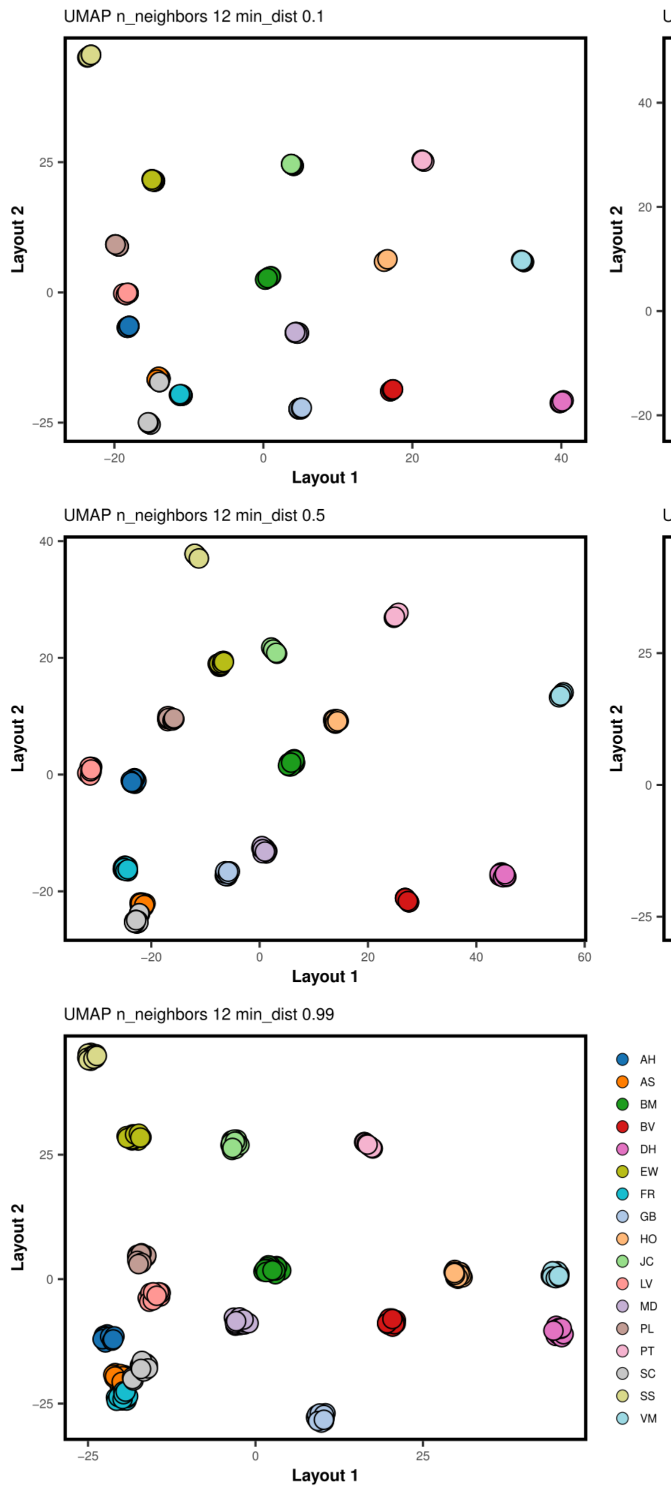
UMAP n_neighbors 12 min_dist 0.1
UMAP n_neighbors 12 min_dist 0.25
UMAP n_neighbors 12 min_dist 0.5
UMAP n_neighbors 12 min_dist 0.8
UMAP n_neighbors 12 min_dist 0.99
Layout 1
Layout 2
AH
AS
BM
BV
DH
EW
FR
GB
HO
JC
LV
MD
PL
PT
SC
SS
VM

**g)**

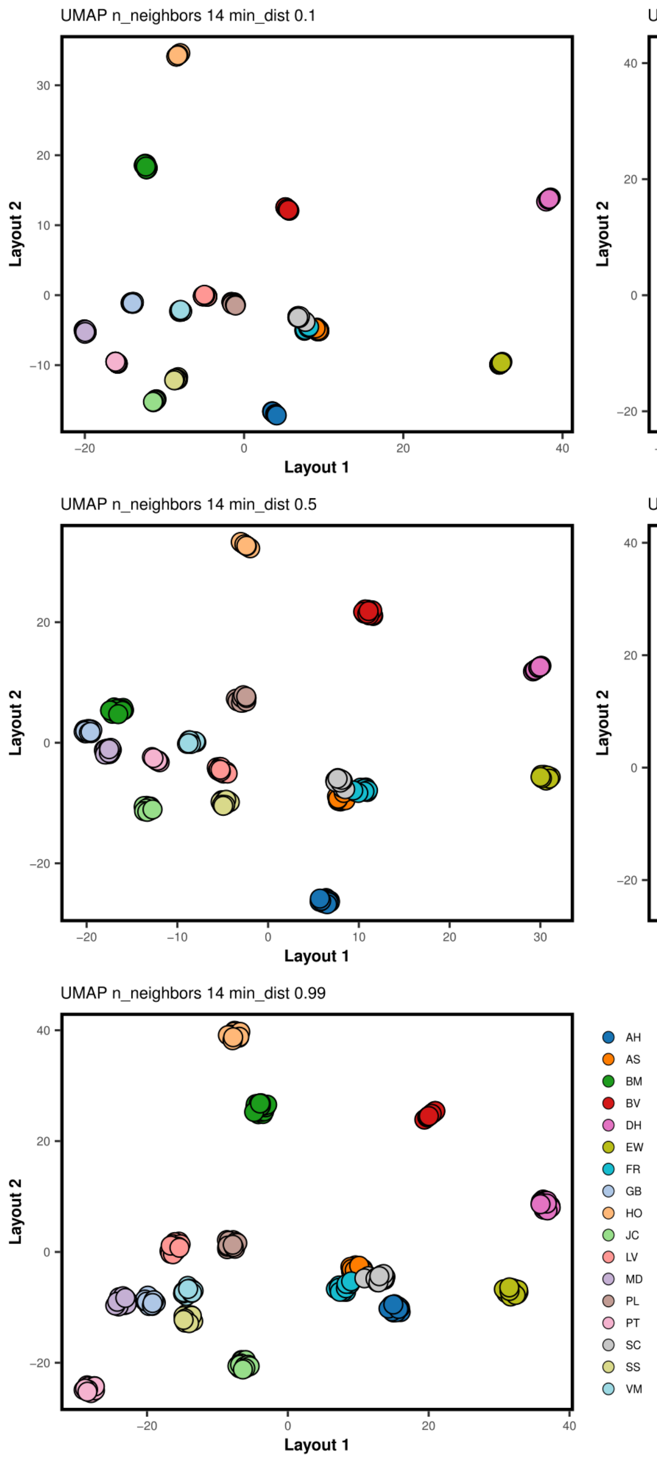
UMAP n_neighbors 14 min_dist 0.1
UMAP n_neighbors 14 min_dist 0.25
UMAP n_neighbors 14 min_dist 0.5
UMAP n_neighbors 14 min_dist 0.8
UMAP n_neighbors 14 min_dist 0.99
Layout 1
Layout 2
AH
AS
BM
BV
DH
EW
FR
GB
HO
JC
LV
MD
PL
PT
SC
SS
VM

**h)**

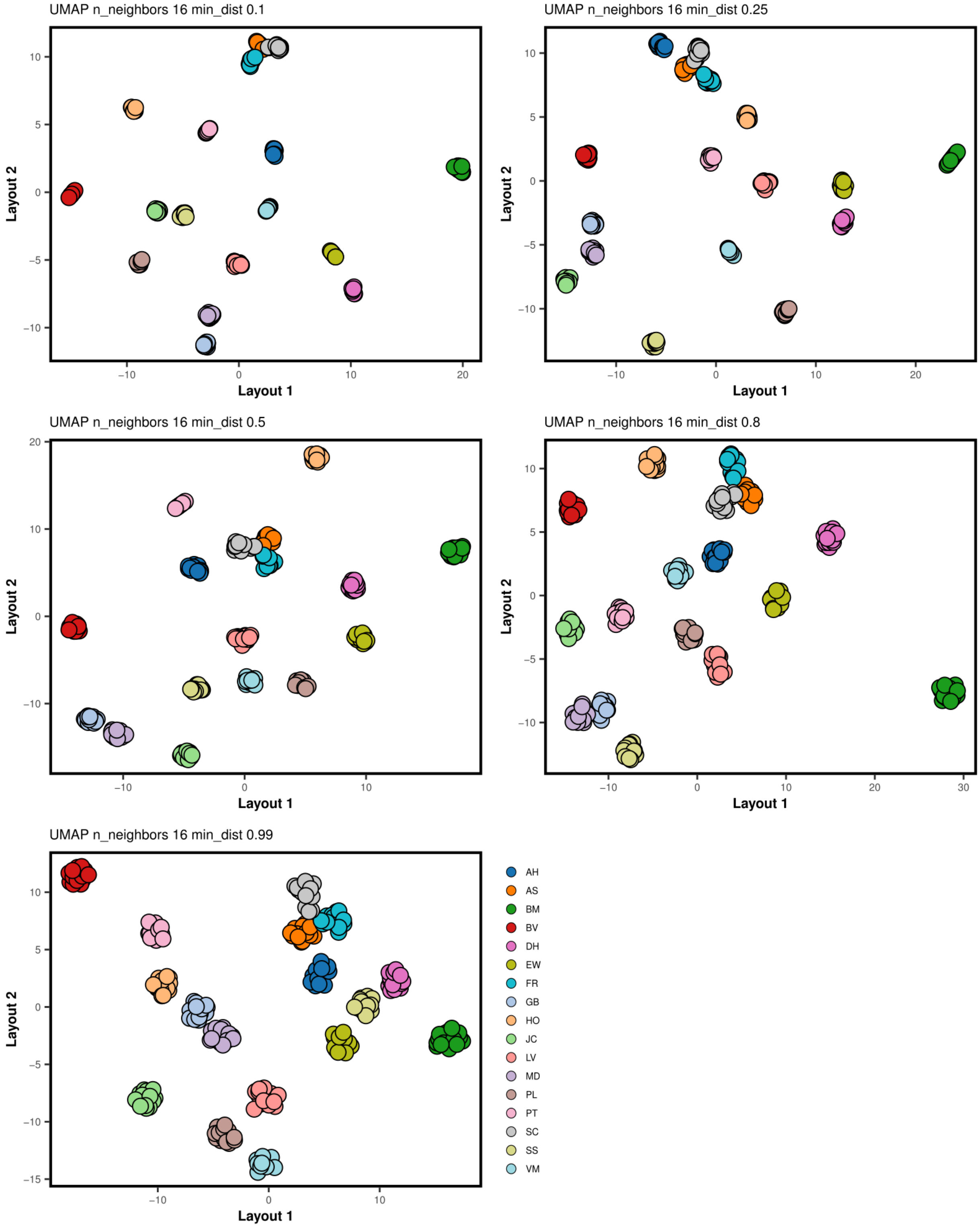
UMAP n_neighbors 16 min_dist 0.1
UMAP n_neighbors 16 min_dist 0.25
UMAP n_neighbors 16 min_dist 0.5
UMAP n_neighbors 16 min_dist 0.8
UMAP n_neighbors 16 min_dist 0.99
Layout 1
Layout 2
AH
AS
BM
BV
DH
EW
FR
GB
HO
JC
LV
MD
PL
PT
SC
SS
VM

**Figure 3.** ADMIXTURE results. **a)** Cross-validation errors values from ADMIXTURE. **b)** ADMIXTURE plots representing estimated membership coefficients for each individual of nine (K= 9), and ten (K= 10) ancestral populations.

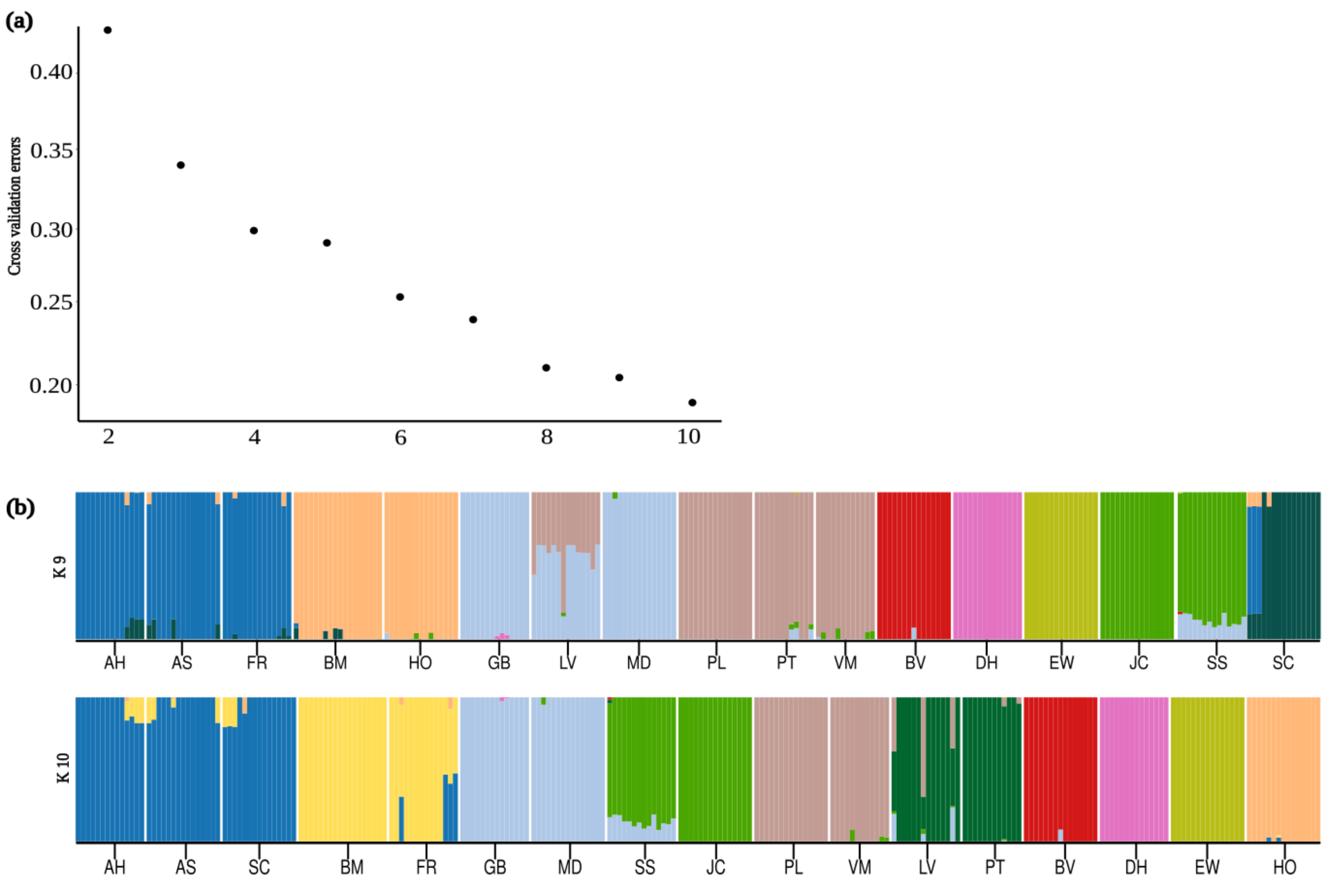
